# Microbiota links to neural dynamics supporting threat processing

**DOI:** 10.1101/2021.04.27.441703

**Authors:** Caitlin V. Hall, Ben J. Harrison, Kartik K. Iyer, Hannah S. Savage, Martha Zakrzewski, Lisa A. Simms, Graham Radford-Smith, Rosalyn J. Moran, Luca Cocchi

## Abstract

There is growing recognition that the composition of the gut microbiota influences behaviour, including responses to threat. The cognitive-interoceptive appraisal of threat-related stimuli relies on dynamic neural computations between the anterior insular (AIC) and the dorsal anterior cingulate (dACC) cortices. If, to what extent, and how microbial consortia influence the activity of this cortical threat processing circuitry is unclear. We addressed this question by combining a threat processing task, neuroimaging, 16S rRNA profiling, and computational modelling in healthy participants. Results showed interactions between high-level ecological indices with threat-related AIC-dACC neural dynamics. At finer taxonomic resolutions, the abundance of *Ruminococcus* was differentially linked to connectivity between, and activity within the AIC and dACC during threat updating. Functional inference analysis provides a strong rationale to motivate future investigations of microbiota-derived metabolites in the observed relationship with threat-related brain processes.

## 1. Introduction

Emerging work suggests that the composition and diversity of the intestinal (gut) microbiota plays a key role in altering brain activity and related behaviour (Cryan and Dinan, 2012). Microbiota-deficient and germ-free mice have provided initial accounts of the effect of microbiota on brain processes, including emotion and affect (Bravo et al., 2011), social behaviour (Sherwin et al., 2019), and cognition (Desbonnet et al., 2015). By extension, there is growing motivation to assess how microbial perturbations are linked to the expression of psychiatric symptoms including the ability to protect against (Clarke et al., 2013), or induce (Neufeld et al., 2011) stress and anxiety-like behaviour.

A key feature underpinning anxiety is the impaired ability to flexibly respond to threat and modify behavioural responses in volatile learning environments (Schiller et al., 2008). Within a broader network of cortico-subcortical regions, the anterior insular cortex (AIC) and dorsal anterior cingulate cortex (dACC) have been the most consistently implicated regions in neuroimaging studies of human threat processing (Fullana et al., 2016). In this capacity, the AIC-dACC circuitry has been hypothesized to support the subjective experience of threat processing via cognitive-interoceptive appraisal mechanisms (Fullana et al., 2016; Harrison et al., 2015; Kalisch and Gerlicher, 2014). Specifically, the AIC is thought to be responsible for generating an awareness of the current emotional and internal physical state (Craig, 2009; Fullana et al., 2016), including gastrointestinal and cardiorespiratory bodily changes (Garfinkel et al., 2015; Rebollo et al., 2018). This information is relayed to the dACC, where its activity has been more directly linked to the cognitive appraisal of bodily sensations of anxiety (Harrison et al., 2015). Given their joint contribution in receiving and processing afferent visceral information, the AIC-dACC network represents a natural candidate to study gut microbiota interactions with higher-level brain function (Kano et al., 2018).

Bidirectional communication between the microbiota and the brain is facilitated by several mechanisms, including immune, endocrine, vagal, and microbiota-derived metabolite signalling (Cryan and Dinan, 2012; Kaelberer et al., 2018; Mayer, 2011; Raybould, 2010). Microbial-derived metabolites like short-chain fatty acids (SCFAs) can activate receptors expressed on the colonic epithelium and within enteroendocrine cells, modulating vagal afferents or immunomodulatory pathways (Koh et al., 2016; Le Poul et al., 2003). Vagal signalling may represent the most direct route through which the microbiota can relay action potentials to cortical regions involved in threat processing (Forsythe et al., 2014; Kaelberer et al., 2018; Raybould, 2010; Rhee et al., 2009). The latency of onset to evoking afferent vagal responses occurs within minutes of intraluminal probiotic administration (Perez-Burgos et al., 2012), while the SCFA butyrate – a by-product of microbial metabolism – has been shown to elicit vagal responses within seconds (Lal et al., 2001). Afferent viscerosensory signals converge in the nucleus tractus solitarius (NTS) and are projected via nuclei in the brainstem – including locus coeruleus, parabrachial nucleus and dorsal raphe nuclei – to higher cortical regions, including the AIC and dACC (Azzalini et al., 2019). The proposed relationship between the viscera and the AIC-dACC network is also supported by the cellular substrate underpinning communication between these two cortical regions: the von Economo neurons (VEN) (Mayer, 2011). These cells contain receptors (serotonin 2b) and peptides (neuromedin B) that are also abundant in the enteric nervous system (Allman et al., 2010). The peculiar expression of receptors and peptides in both viscera and VENs suggest a likely role of the AIC-dACC network in linking brain and gut signals.

Preclinical work has revealed how manipulation of the microbiota impacts threat-related processes (Chu et al., 2019). This work provides preliminary support for the notion that changes in the composition of the microbiota, and associated metabolite production, modulates neural activity in distinct brain regions. However, due to the large variability between the mouse and human microbiota, preclinical observations have not always replicated in human studies (Sherwin et al., 2019). It is also unclear whether human microbiota-brain interactions can be characterised by shifts in high-level ecological measures (e.g., alpha diversity), or whether they emerge within taxonomic scales at finer resolutions. To bridge this knowledge gap, we combined a threat processing paradigm (Savage et al., 2020) (**Fig. 1a**), 16S rRNA gene sequencing (of stool samples), functional magnetic resonance imaging (fMRI), and computational modelling. Using established conditioning procedures, the adopted task facilitates the assessment of both general threat learning as well as threat updating responses in AIC and dACC. Recently, using this task, it was shown that threat updating responses in dACC were especially predictive of subjectively reported bodily anxiety sensations (Savage et al., 2020).

**Fig. 1.**
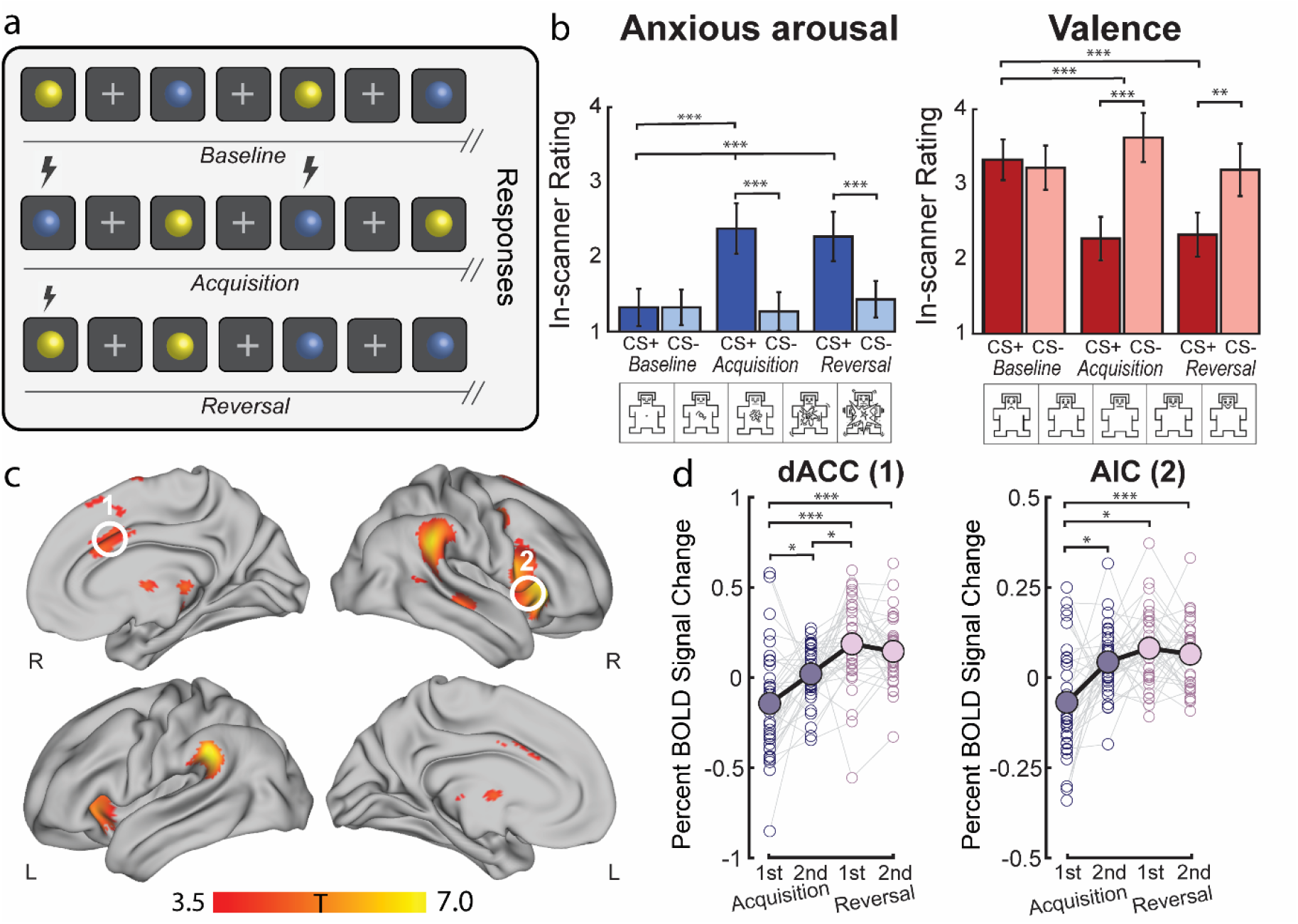
Behavioural results and neural correlates of threat *acquisition* and *reversal*. (**a**) Design of the threat processing paradigm implemented in the MRI scanner. The fMRI task lasted for 17 minutes and had three phases: *baseline* (top row), *acquisition* (middle row), and *reversal* (bottom row). Blue and yellow spheres were used as the conditioned stimuli (CS). Between each CS presentation, a white fixation cross appeared which served as a fixed inter-stimulus interval (ISI). The unconditioned stimulus (US, lightning bolt) was an aversive auditory burst. During *acquisition*, the US co-terminated with one of the CS (forming a threat, CS+) and not with the other (forming a safety, CS-). During *reversal*, the pairing of the US and CS was switched. Immediately after each phase (in-scanner, as a continuation of the task phase), participants were asked to rate the spheres in terms of subjective bodily anxiety sensations and affective valence on a five-point Likert scale (Self-Assessment Manikins, SAM) (Bradley and Lang, 1994). (**b**) Behavioural results for subjective *in-scanner* ratings of the threat and safety signals during *baseline*, *acquisition* and *reversal* for bodily anxiety sensations (left) and affective valence (right). The ratings confirmed the differential aversiveness of the threat relative to the safety signal, and the *acquisition /reversal* compared to the *baseline* stimulus (where no US was present). For post-hoc t-tests (Bonferroni-corrected), ** denotes *p_FWE_* < 0.001 and *** denotes *p_FWE_* < 0.0001. (**c**) The contrast overall (combining *acquisition* and *reversal* task phases) threat (CS+) > safety (CS-) was associated with significant (*p_FWE_* < 0.05 at cluster level, high threshold of p_uncorrected_ < 0.001) group level activation in cortical and subcortical brain regions, including the mid dACC (white circle “1”) and right AIC (white circle “2”) (details in Supplementary Table 1). (**d**) The difference in mean percent BOLD signal change responses between threat and safety signals were assessed in the dACC and AIC during *acquisition* (1^st^ and 2^nd^ half) and *reversal* (1^st^ and 2^nd^ half). Solid dots and black lines represent the group-level mean percent BOLD signal change responses. For post hoc paired t-tests (Bonferroni-corrected), * denotes *p_FWE_* < 0.05, and *** denotes *p_FWE_* < 0.0001.

Our first aim was to extend existing neuroimaging work by demonstrating the effective (causal) connectivity patterns between neural populations within the AIC and the dACC during the assessment of general threat acquisition, and subsequent updating processes. To understand whether inter-individual differences in microbiota covaried with evoked AIC-dACC responses during threat processing, we adopted a Bayesian and multivariate statistical framework (Zeidman et al., 2019). Specifically, effective connectivity measures were assessed against (i) high-level ecological measures (α-diversity and phylum ratio) and (ii) specific (genus-level) measures to discern the resolution/s at which interactions emerge. High-level microbial measures were selected a priori based on previous work (Ley et al., 2006; Lozupone et al., 2012; Mariat et al., 2009). At genus-level, we used a validated data-driven clustering approach (Arumugam et al., 2011) to discern features contributing to the largest variance among samples. The selection of genus-level candidates were consistent with previous literature (Tillisch et al., 2017; Valles-Colomer et al., 2019) and were assessed both independently (multiple regressions) and in the context of a multivariate microbiome-brain relationship (sparse canonical correlation analysis). Finally, guided by preclinical work, we assessed the predicted functional capacity of the microbiota - focusing on pathways involved in SCFA production. This last analysis aimed to provide support to previously proposed causal pathways linking microbiota to brain (Boehme et al., 2019; Lee et al., 2020; van de Wouw et al., 2018).

## 2. Materials and Methods

### 2.1 Participants

The study was approved by the Human Research Ethics Committee of QIMR Berghofer Medical Research Institute (P3435). Written informed consent was obtained for all participants. Thirty-eight healthy adult participants (31.7 ± 8.8 years; 23 female) were recruited from the Brisbane (Australia) metropolitan area by an accredited practising dietitian (APD) (Supplementary Table 1). Exclusion criteria included: a BMI of < 18.5 or > 30.0; current or previous history of a major psychiatric illness or neurological disorder (assessed via neurocognitive assessments performed by a trained psychologist); chronic or clinically significant pulmonary, cardiovascular, gastrointestinal, hepatic, renal, or dermatological functional abnormality as determined by medical history; history of cancer (excluding medically managed squamous or basal cell carcinomas of the skin); history of active, uncontrolled gastrointestinal condition, disease, or irregular bowel movements (including persistent diarrhoea or constipation); history of psoriasis or recurrent eczema; major changes to dietary intake in the past month (self-report); consumption of ≥ 5 standard alcoholic drinks per day; recreational drug use ≥ 1 occasion in the past 3 months; acute disease at time of enrolment; pregnancy or lactation; and use of the following medications within the past 3 months: antibiotics, antifungals, antivirals, antiparasitics, corticosteroids, cytokines, methotrexate or immunosuppressive cytotoxic agents, large doses of commercial probiotics, or selective serotonin reuptake inhibitors. Gastrointestinal and microbiota-related exclusion criteria were adapted from an existing framework provided by The Human Microbiome Project Consortium (Methé et al., 2012). Dietary assessments, including the Traditional Mediterranean diet (TMD) questionnaire, reported gastrointestinal symptoms, current medication use, and recent changes to major food groups were administered by an APD. Each participant completed: i) a health and neurocognitive assessment; ii) home collection of a stool sample, and iii) structural (T1) and functional magnetic resonance (MR) scans. Study requirements were separated into two sessions; the second session completed no longer than 14 days after the first (4.8 ± 3.9 days between sessions, mean ± standard deviation).

### 2.2 Experimental Paradigm

A validated framework to study threat learning and updating is a Pavlovian threat processing paradigm, wherein an emotionally neutral stimulus (CS) is differentially conditioned with an aversive auditory event (US) before the contingency is subsequently switched in a later *reversal* phase. The fMRI task, previously used by Savage et al. (Savage et al., 2020), lasted for 17 minutes and had three phases: *baseline*, *acquisition*, and *reversal* (**Fig. 1a**). Blue and yellow spheres, presented for two seconds against a black background, were used as the conditioned stimuli (CS). Between each CS presentation, a white fixation cross appeared, which served as a fixed inter-stimulus interval (ISI, 12 seconds). The unconditioned stimulus (US) was an aversive auditory (white noise) burst (50ms) presented at 75-100dB that occurred at the end of CS+ presentation and co-terminated with the CS. The US volume was determined during a pre-task calibration, where participants were asked to rate the unpleasantness/averseness of the white noise. The white noise burst serving as the US has been validated in previous fMRI studies (Harrison et al., 2017; Savage et al., 2020). During *baseline*, the CS were each presented 5 times and no US occurred. During *acquisition*, the US co-terminated with one of the CS (forming a CS+) and not with the other (forming a CS-, safety). The color of the CS+ was counterbalanced across subjects and the CS-US pairing occurred one third of the time, enabling the classification of CS+unpaired trials and the subsequent analysis of threat responses without US confounding. During the reversal phase, the pairing of the US and CS was switched. 10 presentations of the CS+ unpaired, 5 of the CS+ paired and 10 presentations of the CS- occurred during both acquisition and reversal task phases, with no more than two consecutive trials of the same stimuli. Throughout the paper, the CS+unpaired is simply referred to as threat (initial threat for the acquisition phase and updated threat for the reversal phase) for ease of readability. Immediately after each phase (in-scanner, as a continuation of the task phase), participants were asked to rate the spheres in terms of anticipatory anxiety and emotional valence on a five-point Likert scale (Self-Assessment Manikins, SAM) (Bradley and Lang, 1994). Upon completion of each task phase, participants were instructed to respond to questions assessing subjective anticipatory anxiety (“How anxious did the [blue or yellow] sphere make you feel?” Responses ranged from 1 = ‘not at all anxious’ to 5 = ‘very anxious’); and emotional valence (“How unpleasant/pleasant did you find the [blue or yellow] sphere?” Responses ranged from 1 = ‘very unpleasant’ to 5 = ‘very pleasant’). Responses were made using a hand-held button-press box held in the participant’s dominant hand. Prior to the scan, participants were familiarized with the scales, response box, and the volume of the US.

### 2.3 Image acquisition and pre-processing

Images were acquired on a 3T Siemens Prisma MR scanner equipped with a 64-channel head coil. For the fMRI task, whole brain echo-planar images were acquired using a multiband sequence (multiband factor of 8, GRAPPA factor of 1). 1227 volumes were collected with the following parameters: voxel size = 2mm^3^; TR = 810ms; TE = 30mm; flip angle = 53°; FOV = 212mm; slice thickness = 2mm; 72 slices; 0.63ms echo spacing. T1 and spin echo (anterior to posterior and posterior to anterior directions) images were also acquired to assist with pre-processing of the functional data. Image pre-processing was performed using fMRIPrep version 1.3.2(Esteban et al., 2019) and Python scripts developed in-house (available online: https://zenodo.org/record/3556980#.XlZUr6j7TIU). Briefly, pre-processing involved head-motion correction, susceptibility distortion correction, confounds estimation, coregistration, and regression of nuisance covariates including WM, CSF, and the six head motion parameters. Smoothing using a FWHM Gaussian filter (10mm) was performed with the SPM12 software (Wellcome Trust Centre for Neuroimaging, UK).

### 2.4 Stool collection

Participants were provided with a stool collection kit customized for this study and were advised to collect and return the stool sample within 24-48 hours before/after the MR scan. The kit included a stool nucleic acid collection and preservation tube (Norgen Biotek Corp., Thorold, Ontario, Canada), written instructions on how and when to collect the stool sample, and a pair of plastic gloves. Upon return, each stool sample was labelled with a de-identified participant code, and transported and stored in a −80°C freezer until sample processing.

### 2.5 DNA preparation and 16S rRNA gene sequencing

Tissue homogenization was performed using tubes containing 1.4mm ceramic beads (Precellys Lysing Kit). DNA was extracted from samples and quantitated using Nanodrop 2000 (Thermo Scientific). PCR amplification was performed on the V3-V4 hypervariable region of the 16S rRNA gene, and sequenced on a MiSeq sequencer (Australian Genome Research Facility, Brisbane). Paired-end reads were joined using PEAR v0.9.6 and PCR primer sequences were removed using Cudadapt. Sequence data were processed using Quantitative Insights Into Microbial Ecology (QIIME) software suite v1.9.1 using default settings. USEARCH v8.0 was used to cluster the sequences into Operational Taxonomic Units (OTUs) using the identity threshold of 97%. Only OTUs with at least two reads were included. Representative sequences of each OTU were taxonomically classified using USEARCH, and aligned to the Greengenes reference alignment (v13.8) using PyNAST. The OTU table was normalized (total sum scaling) and square root transformed to account for the non-normal distribution of taxonomic count data. Samples were rarefied to a read depth of 5,511 for diversity analyses. For α-diversity indices, we calculated Chao1, Shannon, Simpson, and inverse Simpson measures. For our high level assessment linking microbiota to brain, we opted to use Inverse Simpson diversity due to its large inter-individual variability. The functional capacity of each sample was predicted using a computational modelling approach, called Phylogenetic Investigation of Communities by Reconstruction of Unobserved States (PICRUSt) (Langille et al., 2013). Gene counts encoding enzymes were then predicted using the *metagenome_predictions.py* script which were then mapped onto Kyoto Encyclopaedia of Genes and Genomes (KEGG) pathways (Kanehisa and Goto, 2000). We focused specifically on gene counts encoding terminal enzymes (i.e., final conversions) involved in SCFA production, identified using the KEGG reference pathways for butanoate metabolism (KEGG ref: 00650) and propanoate metabolism (KEGG ref: 00640). Gene counts that were not represented in at least 50% of samples were excluded from further analyses. The contributions of OTUs were predicted using the *metagenome_contributions.py* script, and were further summarized into percent contributions at the genus, order, and phylum level. Tutorial steps outlining the full PICRUSt pipeline can be found online (http://picrust.github.io/picrust/).

### 2.6 Neuroimaging analyses

For each participant, the pre-processed images were included in a first-level GLM analysis, performed with SPM12. For all three phases, the onsets of each CS event-type (*baseline*: n = 5 per CS; *acquisition* and *reversal*: CS-(safety) = 10, CS+unpaired (threat) n = 10, CS+paired, n = 5) were modelled as a series of delta (stick) functions, convoluted to the canonical haemodynamic response function (HRF). Model parameters were estimated using Restricted Maximum Likelihood (ReML). The resulting set of first-level contrast images were carried forward to group-level random-effects analysis. The difference in percent BOLD signal change between the threat and safety signals was extracted from the AIC and dACC during the *acquisition* and *reversal* task phases using the Marsbar toolbox for SPM (Brett et al., 2002). To disentangle the effects within each phase, we independently examined the differences between early (first 5 CS+ unpaired presentations) and late (last 5 CS+ unpaired presentations) mean responses in both *the acquisition* and *reversal* task phases. Neuroimaging analyses were reproduced using an alternative smoothing kernel of 5mm and demonstrated consistent group-level GLM and DCM results (Supplementary Figure 7, Supplementary Table 2).

### 2.7 Specification and inversion of DCMs at the first level

We used DCM, a computational framework to investigate the effective (directed) connectivity between and within (self-connections) cortical regions. To do this, DCM for fMRI couples a bilinear model of neural dynamics with a biophysical model of hemodynamics. Details regarding this method can be found elsewhere (Friston et al., 2003). Subject-specific regions of interest (ROIs) were defined as a 5mm sphere centred over each subject’s peak functional activation, constrained within the functional mask generated by the group-level contrast overall (*acquisition* and *reversal*) threat > safety (*p*_FWE_ < 0.05 at cluster-level, high threshold of p_uncorr_ < 0.001; **Fig. 1c**). Subjects who had no or minimal task-related fMRI activation in AIC and dACC were excluded from the DCM analysis (p_uncorr_ < 0.1) (Supplementary Figures 8 and 9). Having identified the AIC (x=32, y=34, z=0) and dACC (x=2, y=20, z=28) coordinates at the group level, the spatially closest (Euclidian distance) peak coordinates for each individual subject were manually located and the time series were extracted as the peak eigenvariate for all remaining participants (n = 33, exclusion of 5). Inspection of the ROIs showed that they were all located within 10mm from the group level peak (Supplementary Figure 1) and were anatomically consistent with previous work (Savage et al., 2020; Tian and Zalesky, 2018). DCM, as a hypothesis-driven framework, operates on a user-defined model space specified through the choice of: (i) endogenous (context-independent average) connections, (ii) contextual (experimental) modulation of endogenous connections, and (iii) direct inputs (e.g., CS stimuli) to the system. We constructed three models, all of which considered bidirectional endogenous connections between the AIC and dACC, and intrinsic connections at both brain regions (i.e., a fully-connected A-matrix). The driving inputs (C-matrix) consisted of visual (threat/safety) and auditory (CS+paired) stimulus, modelling the possibility that input could enter at either the AIC or dACC. This allowed the US (white noise) to affect driving inputs separately from the CS+ for which there was a threat response but no auditory sound. The difference between models arises from the choice of contextual modulators (B-matrix). Specifically, the B-matrix was specified to test whether there were differential modulatory effects between the two task phases: *acquisition* and *reversal*. The first model accounted for the modulation evoked exclusively by the CS+ events during *acquisition* and *reversal* (each phase considered as a separate modulator, **Fig. 2b,** model 1). The second model accounts for the modulatory effects of all events (all trials of CS+, safety and ISIs) acquired during the *acquisition* and *reversal* task phases (i.e. to model tonic block effects from threat processing; **Fig. 2b,** model 2). Finally, the third model combines the two aforementioned models, accounting for both threat-related, and block-related events for *acquisition* and *reversal task phases* (**Fig. 2b,** model 3). Model inversion was performed for each subject using the DCM12 routines implemented in SPM12. Bayesian model selection with random effects analysis was used to select the most likely model given the data (as assessed by the highest exceedance probability).

**Fig. 2.**
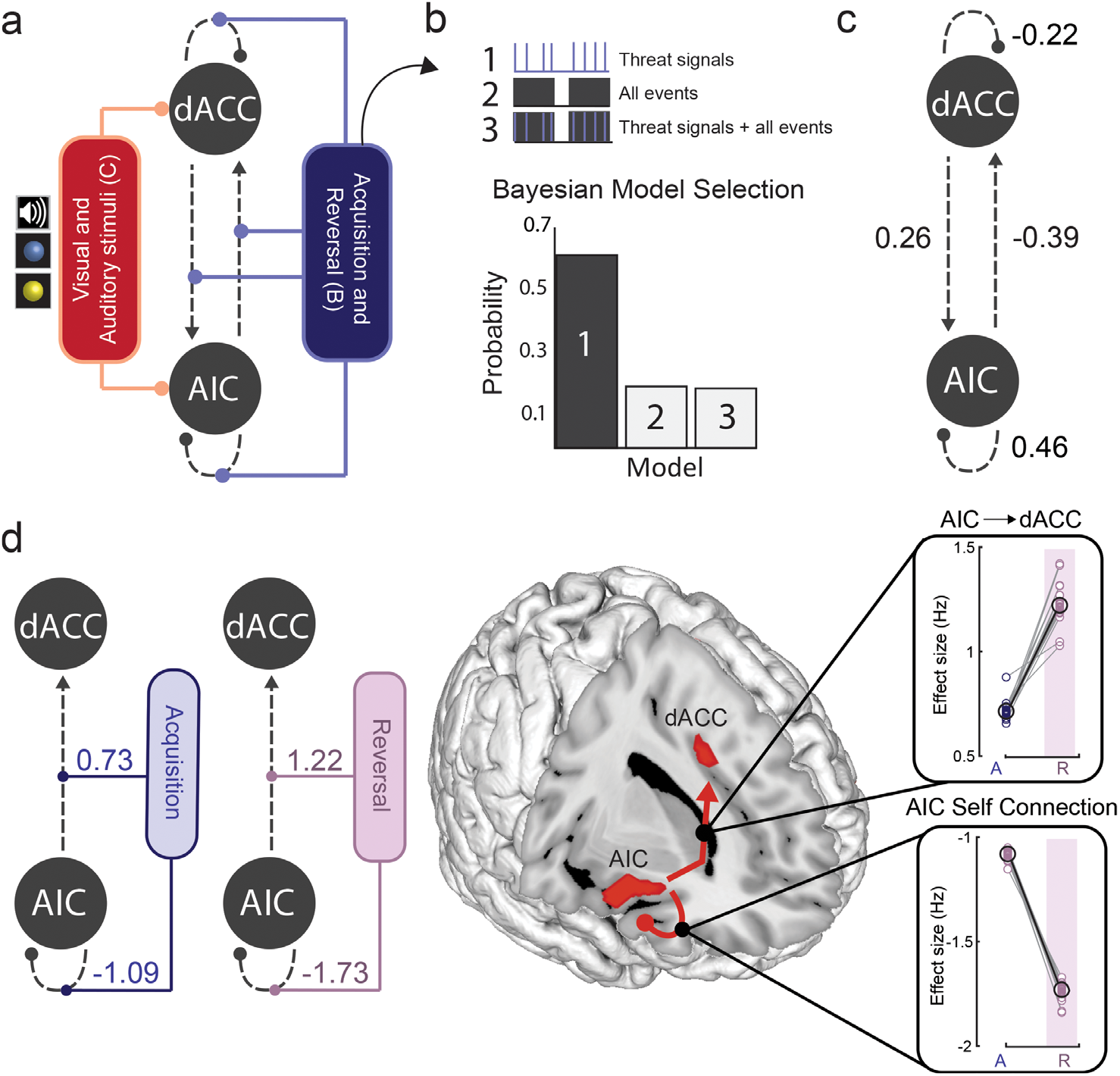
Neural dynamics supporting threat *learning* and *updating*. (**a**) Specification of the DCM model space in terms of: (i) task-independent effective connectivity (grey, dashed lines) (A-matrix); (ii) modulatory connections (B-matrix) (blue), including threat signals in both *acquisition* and *reversal* task phases; and (iii) direct inputs to the system (C-matrix) comprising visual (all CS events) and auditory (US) stimuli (red). (**b**) Three models were estimated for each subject (see text for details). The difference between models arises from the specification of contextual modulators (threat signals, all trials, or both). Bayesian Model Selection showed that the exclusive modulation by threat signals (model 1) best explained the fMRI data (as accessed by the highest exceedance probability). (**c**) Results from Bayesian Model Reduction (BMR) on second level Parametric Empirical Bayes analysis of trial-independent (fixed) connections across individuals. Results showed a positive modulation from the dACC to the AIC, a negative modulation from the AIC to the dACC, and local effects within both regions. (**d**) BMR results showed a significant modulatory effect of threat signals on patterns of effective connectivity during *acquisition* and *reversal* (left). Results highlight very consistent modulatory effects of task phase (from *acquisition* to *reversal*) on AIC ƒ dACC (top, right) and AIC self-connections (bottom, right).

### 2.8 Second-level Parametric Empirical Bayes analysis

The winning DCM model was brought forward to a second-level Parametric Empirical Bayes analysis to investigate where the mean endogenous (task-independent) and modulatory effects were expressed, as well as assessing inter-subject, microbiome-associated variability in responses. To do this, the PEB scheme begins by collating the estimated parameters of interest from all subjects, including the expected values of the parameters, their covariance matrices and approximate likelihoods. Next, Bayesian model comparison was performed using these first level model parameters to infer group effects. The group mean was modelled by including a constant term in the design matrix. Bayesian model reduction over the second level parameters was subsequently performed, which performs a greedy search over all parameters, and prunes away those that do not contribute to the model’s log evidence (Friston et al., 2016). With an understanding of the group-level effects, we then performed a second PEB, this time specifying a design matrix with three regressors. The first regressor modelled the group mean, and the second and third modelled the microbiota covariates (orthogonalized and mean-centred). High-level microbial measures including the ratio between Bacteroidetes/Firmicutes (B/F), and α-diversity - were selected a priori based on previous work (Ley et al., 2006; Lozupone et al., 2012; Mariat et al., 2009). We note that unlike previous work (Ley et al., 2006), we did not observe a significant relationship between B/F ratio and body mass index (BMI) (R = −0.21, p=0.25). PEB returns each parameter, where covariate-specific and group means are reported in terms of the expected values (Ep) and their corresponding posterior probabilities (Pp). Pp > 0.95 were considered to have a non-zero effect.

### 2.9 Enterotype clustering

To perform the brain-microbiota assessment at genus-level, we reduced the dimensionality of the microbiota into features that represented the greatest sources of variability in the data. Microbiota samples were clustered into enterotypes using methods previously described in Arumugam et al. (Arumugam et al., 2011), which are readily available online (http://enterotype.embl.de/enterotypes.html). We used the Calinski-Harabasz (CH) index to identify the optimal number of clusters (from *k* = 2 to *k* = 20). The silhouette index (SI) was adopted to assess how similar a sample was to its own cluster (cohesion), compared to other clusters (range of −1 to +1, where a high positive value indicates that samples are well aligned to its own cluster).

### 2.10 Co-abundance network construction

Using the relative abundances at genus-level (total sum scaled and square root transformed), we computed the Pearson correlation coefficients between the four enterotype-driving genera (*Bacteroides, Ruminococcus, Oscillospira,* and *Prevotella*) and all other genera to generate an undirected, weighted co-occurrence network. Positive (co-occurrence) and negative (co-exclusion or competitive) relationships, using cut-offs at > 0.39 and < −0.39 respectively, were visualized using the interactive platform Gephi (Version 0.9.2), using the force atlas template (Bastian et al., 2009). To confirm the robustness of network interactions, we also computed *Sparcc* correlations (Friedman and Alm, 2012) on count data collapsed at genus level (Supplementary Figure 6).

### 2.11 Multiple regression analysis

Multiple linear regression analysis was used to test the association between AIC-dACC responses with the relative abundance of each driving genera. To avoid overfitting the models, we used PCA to reduce the dimensionality of each of the normalized (z-scored) sets of brain measures. For PCA inputs, we used the DCM modulatory connections for threat *conditioning* (4 parameters) and *reversal* (4 parameters), as well as the percent signal change values for both task phases in the AIC (4 parameters) and dACC (4 parameters). If the first PCA explained < 50% variance, the 2^nd^ PCA component was also included. This resulted in 7 PCA components (2 × threat *conditioning*; 1 × threat *reversal*; 2 × AIC percent BOLD signal change; 2 × dACC percent BOLD signal change) representing our 16 brain measures (Supplementary Figure 4).

### 2.12 Sparse Canonical Correlation Analysis

*l*_1_-norm regularized sparse canonical correlation analysis (sCCA) was implemented using default settings in the R package, “PMA” (Witten et al., 2009). This approach has been designed to partially alleviate modelling challenges in small sample sizes by introducing a penalty for the elements of the weight vectors (Wang et al., 2020). The aim of this analysis is to reduce the variable set according to their most important directions of linear variation, while allowing the microbiota-brain associations to be interpreted within the original variable space. In this implementation, the tuning parameter was automatically selected for each dataset (microbiota and brain) using a permutation scheme (n = 10,000), repeated across ten different candidate sparsity parameters (ranging from 0.1 – 0.7). The best penalty for each dataset was selected based on the highest z-statistic, and the sCCA was then repeated using this parameter (Supplementary Figure 5a). Leave-one-out (LOO) cross-validation was performed to confirm the absence of any single subject outliers (Supplementary Figure 5b). A secondary cross-validation was performed by randomly removing 15% of the total sample (permutations = 1000) and assessing the stability of the (a) sCCA correlation and (b) brain and microbiota weights (Supplementary Figure 5b). An overview of the complete pre-processing and analysis pipeline is shown in Supplementary Figure 10.

## 3. Results

### 3.1 Behavioural Results

Participants completed a differential threat conditioning task that involved initial learning (‘*acquisition* phase’) of threat (CS+) and safety (CS-) signal associations, and subsequent updating of these associations during a ‘*reversal* phase’ (Savage et al., 2020) (**Fig. 1a**). At the end of each task phase (including a baseline phase), participants rated the extent to which the threat and safety signals evoked bodily anxiety sensations, respectively, as well as their affective valence. Consistent with previous work (Savage et al., 2020), results from the participants’ subjective in-scanner ratings confirmed that the threat signals induced significantly (p < 0.001) higher bodily anxiety sensations (anxious arousal) and were more unpleasant (negative valence) than the safety signals during both *acquisition* and *reversal* phases, as well as compared to the threat signals during *baseline* (pre-conditioning, where no US was present) (**Fig. 1b**, Supplementary Note 1). As expected, there were no significant differences in participants’ subjective ratings of the threat and safety signals at *baseline* (**Fig. 1b,** Supplementary Note 1). Results from two-factor repeated measures ANOVAs and post-hoc paired t-tests are reported in Supplementary Note 1.

### 3.2 Computations within specialized brain regions

As participants demonstrated successful differential learning across both the *acquisition* and *reversal* task phases, we examined the general neural correlates of threat learning by averaging responses to the threat and safety signals across these phases. That is, we assessed brain regions showing higher activity to the threat signals during *acquisition* and *reversal,* when compared to the safety signal. Results showed a robust group-level difference in the right AIC and mid dACC (*p*_FWE_ < 0.05 at cluster-level, cluster isolated using *p*_uncorr_ < 0.001, **Fig. 1c**). Additional brain regions, including the middle frontal gyrus, and ventral striatum were also more active in the overall (initial and reversed) threat > safety contrast (Supplementary Table 3). In line with our core aim, and existing evidence supporting their role in recruiting cognitive-interoceptive appraisal mechanisms, the right AIC and mid dACC were selected as our regions of interest (Supplementary Figure 1). While this contrast reflects the joint neural correlates of *acquisition* and *reversal* phases, we also examined the specificity of AIC and dACC activation patterns over time (comparing *acquisition* to *reversal*). To do this, we extracted the difference in mean fMRI percent signal change responses between threat and safety signals relative to the baseline of the AIC and dACC (regions defined by two 5mm spheres using the peak maxima from the overall contrast) (Brett et al., 2002). In line with previous work using a similar task (Schiller et al., 2008), we separately assessed brain responses during the early (1^st^ half) and late (2^nd^ half) *acquisition* and *reversal* phases to examine putative learning-related changes (Brett et al., 2002). Results of a one-way repeated-measures ANOVA showed a global difference across conditions for both dACC (F_3, 99_ = 13.99, *p* = 1.12 × 10^−7^, η_p_^2^ = 0.24) and AIC (F_3, 99_ = 3.01, *p* = 0.03, η_p_^2^ = 0.07) (**Fig. 1d).** In the dACC, post hoc paired t-tests (Bonferroni-corrected) showed a significant increase in response from early *acquisition* to late *acquisition* (t_33_ = −2.89, *p_FWE_* = 0.04), and from *acquisition* to *reversal* (early *acquisition* vs early *reversal*, t_33_ = −4.63, *p_FWE_* = 3.34 × 10^−4^; early *acquisition* vs late *reversal*, t_33_ = −4.84, *p_FWE_* = 1.75 × 10^−4^; late *acquisition* vs early *reversal*, t_33_ = −3.27, *p_FWE_* = 0.02) (**Fig. 1d**). In the AIC, an increase in response was observed in the transition from early *acquisition* to late *acquisition* (t_33_ = −3.43, *p_FWE_* = 0.01) and early *acquisition* to *reversal* (early *acquisition* vs early *reversal*, t_33_ = −4.14, p_*FWE*_ = 0.001; early *acquisition* to late *reversal,* t_33_ = −4.46, *p_FWE_* = 5.30 × 10^−4^) (**Fig. 1d**). Stronger shifts in dACC and AIC responses were captured between task phases, rather than between early and late divisions within each phase.

### 3.3 Neural dynamics supporting threat processing

We used Dynamic Causal Modelling (DCM, see methods) to study neural interactions between the AIC and dACC. We estimated three model variants for each subject, assuming bidirectional connections, modulatory effects on all possible connections, and allowing direct inputs at both nodes (**Fig. 2a**). The three models occupied distinct functional pathways that could be altered by threat and/or all task-related events (**Fig. 2a-b**). More specifically, the first model tested for modulatory effects evoked exclusively by the threat signals during *acquisition* and *reversal* (two regressors, as each phase was considered as a separate modulator, **Fig. 2b**, model 1). The second model accounted for the modulatory effects of *all* trials occurring during the *acquisition* and *reversal* phases, including threat, safety, and inter-stimulus intervals (two regressors, **Fig. 2b**, model 2). The third model combined the two aforementioned models, accounting for both threat-related, and trial-related events for *acquisition* and *reversal* (four regressors, **Fig. 2b,** model 3). This approach allowed us to assess whether directed connectivity strengths within the AIC-dACC network were modulated exclusively by threat-related stimuli, or whether a ‘tonic-like’ effect exists (responses to threat produces a constant or increasing modulatory response over the duration of the task).

Bayesian model selection with random effects indicated the first model (threat-based signals) as the most likely given the data (as assessed by exceedance probability) (**Fig. 2b**). To identify significant connections within this model at the group-level, we applied Parametric Empirical Bayes (PEB). This uses Bayesian model reduction (BMR) to automatically search over all parameters and prune effects that do not meaningfully contribute to the model’s log evidence. Significant effects are here defined by parameters where the posterior probability (Pp) of a non-zero effect was ≥ 0.95. We started by assessing connection strengths independent of task activation. These results showed that across all trials there was a positive connection from the dACC to the AIC, an inhibitory connection from the AIC to the dACC, and local responses at the AIC and dACC that were consistent across individuals (**Fig. 2c**). We next tested the effects of threat *acquisition* and *reversal* on all potential connections (AIC to dACC, dACC to AIC, AIC and dACC self-connections). In the *acquisition* phase, the threat-based signals enhanced bottom-up connectivity from the AIC to dACC, as well as local inhibitory responses at the AIC (**Fig. 2d**, blue). The same pattern of results was observed during threat *reversal,* but with stronger effects overall (**Fig. 2d**, pink). The expected values (Ep) and Pps for all parameters are presented in Supplementary Table 4.

### 3.4 α-diversity covaries with AIC-dACC dynamics

Addressing the first of our microbiota-brain aims, we investigated whether patterns of effective (modulatory) connections underpinning threat learning and updating covaried with high-level microbial properties. Emerging evidence suggests that high-level microbial measures, including the *Bacteroidetes* to *Firmicutes* (B/F) ratio and α-diversity (a joint measure of the richness and evenness of microorganisms), are important and distinct indicators of the promotion and maintenance of host health (Ley et al., 2006; Lozupone et al., 2012; Mariat et al., 2009). As there was sufficient variability in these measures across individuals, the B/F ratio and α-diversity constituted two group-level parametric covariates (**Fig. 3a-b**). To test for an effect of these covariates on directed patterns of connectivity, we again applied the Parametric Empirical Bayes method - this time testing for the presence of microbial covariation within the neural data. We assessed different combinations of covariates: group-level effects, a single microbiota covariate, or both microbiota covariates (**Fig. 3c**). Results indicated that the most likely model included the second microbiota covariate, α-diversity (model 3, posterior probability of 0.96, **Fig. 3c**). To confirm the validity of these findings, we repeated the analyses on surrogate models generated by permuting values within the microbiota covariates. The posterior probability of our winning model was significantly higher compared to the random surrogate models (n = 1000 permutations) (**Fig. 3d**). To isolate the significant parameters contributing to the model evidence, i.e. specific connections that covaried with α-diversity, we applied BMR on the winning model (**Fig. 3e**). Inter-individual measures of α-diversity covaried with regulatory (inhibitory) control from the dACC to AIC, and local inhibitory responses at the AIC during threat *reversal* (**Fig. 3e-f**). A confirmatory analysis further suggested that stronger signalling from the dACC to the AIC during *reversal* downregulates AIC activity (inhibitory modulation) (Supplementary Note 2). To control for common confounds often associated with human microbiota-based research, we adopted the relevant exclusion criteria from The Human Microbiome Project Consortium (Methé et al., 2012) and ensured that faecal samples were collected within a 48-hour window before or after the scanning session (see Methods). Furthermore, a confirmatory analysis controlling for the effects of sex and age showed highly consistent results.

**Fig. 3.**
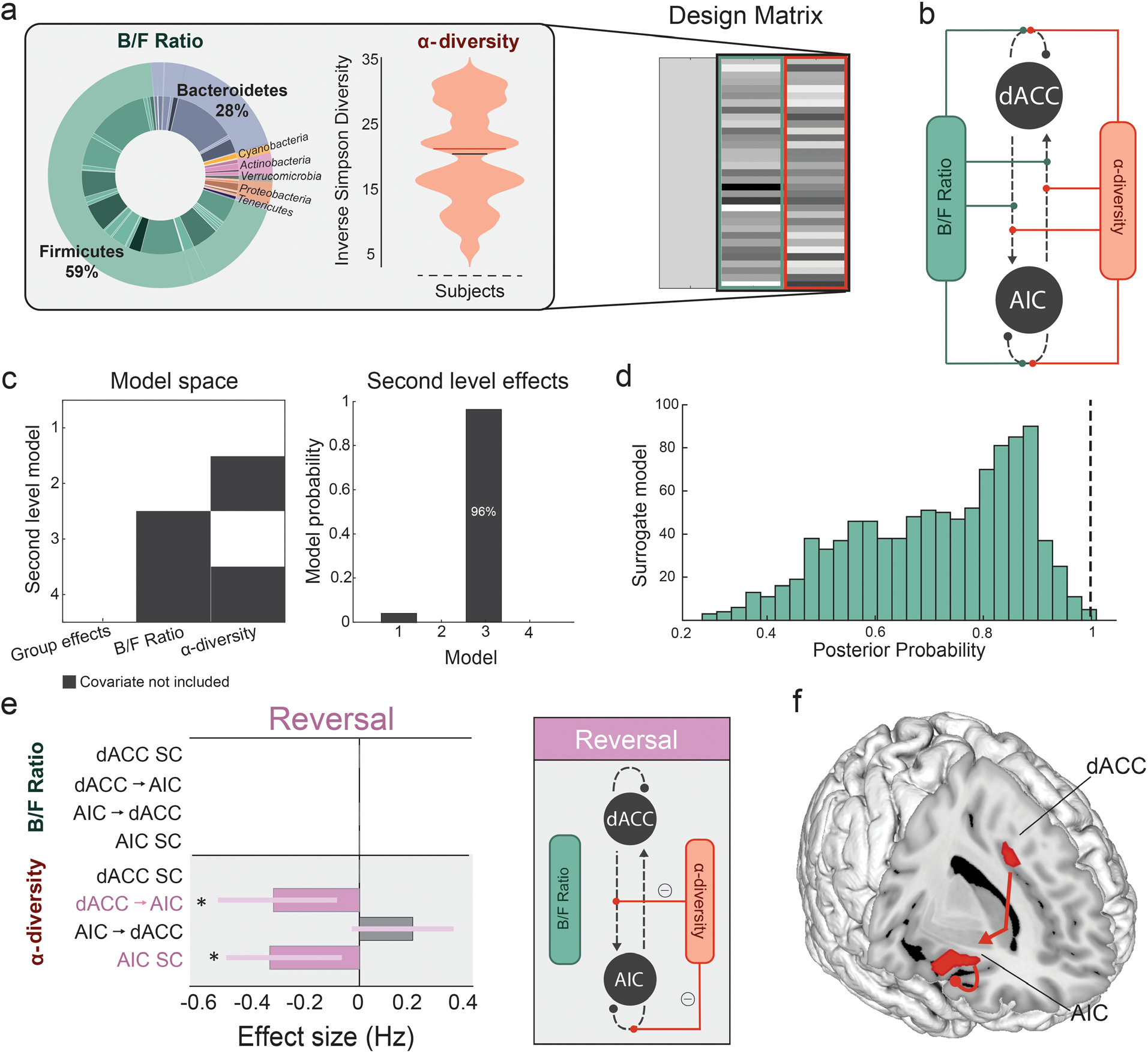
Effects of high-level microbial properties on threat *learning* and *updating*. (**a**) Inlet (left) shows the mean proportions of B/F, and the violin plot (middle) shows the distribution of α-diversity (Inverse Simpson diversity) scores in our samples (n = 38). These two microbiota features constituted our two regressors in the Parametric Empirical Bayes model (design matrix columns two and three, right). The first column models the group mean. (**b**) Specification of the model space showing all possible modulatory connections where the microbiota features can interact within the AIC-dACC network. (**c**) Model space (left) showing possible second level models (including a null model), where both covariates (model 1), one covariate (model 2 and 3), or no covariates (model 4, null) contribute to the model evidence. The winning second level model (right) included the second covariate (model 3), at a posterior probability (Pp) of 0.96. (**d**) Distribution of Pp results from surrogate testing. Dashed black line indicates the Pp (0.96) of the winning model for the original (non-permuted) data. (**e**) Results from Bayesian Model Reduction (BMR) showing the effect sizes (expressed in Hz) of modulatory connections associated with α-diversity during threat *reversal*. Significant parameters are those with a Pp > 0.95, indicated with an asterisk. The length of the bars corresponds to the expected probability (Ep) and the error bars are 90% Bayesian confidence intervals. SC represents self-connections. (**f**) Anatomical representation showing the significant modulatory connections associated with α-diversity during the threat *reversal* phase.

### 3.5 The human microbiota exhibits variability of enterotype-driving genera

We next investigated whether similar patterns of microbiota-brain interactions emerged at lower levels of the taxonomic hierarchy (i.e., from coarse-level measures to genus level). To reduce the dimensionality of the microbiota into features that represent the greatest sources of variability in the data, we performed an enterotype analysis which uses the Partitioning Around Medoids (PAM) (Kaufman and Rousseeuw, 2009) clustering (Supplementary Note 3; Supplementary Figure 2). In line with previous work (Arumugam et al., 2011), between-class analysis revealed that clusters were driven by inter-individual variability in the following genera (Falony et al., 2016): *Bacteroides* (enterotype 1), *Ruminococcus*/*Oscillospira* (enterotype 2), and *Prevotella* (enterotype 3) (Supplementary Figure 2). While clusters were non-random (Supplementary Note 3; Supplementary Figure 3), they were also not exclusively discrete. Instead, visualization of the samples suggested that the data were distributed along a continuum (Supplementary Figure 2). Based on these findings, our subsequent analyses focused on a dimensional assessment of the link between the four discriminate genera and brain dynamics. This approach was also motivated by previous work, suggesting an interrogation of dominant taxa that drive separation between samples, rather than the enterotype classifications themselves, as features to link with clinical or behavioural variables (Cheng and Ning, 2019; Gorvitovskaia et al., 2016).

### 3.6 Distinct effects of genus abundance on threat-processing brain dynamics

We first assessed whether AIC-dACC network strengths were linearly correlated with the abundance of each driving genus independently. Specifically, separate multiple linear regressions were used to assess the association between the abundance of each candidate genus (*Bacteroides, Oscillospira, Ruminococcus,* and *Prevotella*) with distinct brain indices (threat-induced connectivity strengths and percent BOLD signal change). To reduce the dimensionality and improve interpretability, a principal components analysis (PCA) was separately applied to the values of (a) AIC-dACC connectivity during threat *acquisition*; (b) AIC-dACC connectivity during threat *reversal*; and percent BOLD signal change responses in the (c) AIC; and (d) dACC. When the first principal component (PC) for each brain set (a-d) explained < 50% variance, the second component was also included. This resulted in 7 PCs (2 × threat *acquisition*, 1 × threat *reversal*, 2 × AIC, and 2 × dACC percent signal change) which together constituted our regression features (Supplementary Figure 4). PCs representing brain variables explained a significant amount of the variance in *Ruminococcus* (R^2^ = 0.49, F_(25, 33)_= 3.48, *p* = 0.01, **Fig. 4a**). *Ruminococcus* abundance yielded three significant regression weights (β = −0.47, t_(25)_ = −2.65, *p* = 0.01; β = −0.37, t_(25)_ = −2.43, *p =* 0.02; β = 0.45, t_(25)_ = 2.71, *p* = 0.01), supporting an interaction with connectivity responses during threat *acquisition* (1^st^ PCA for threat *acquisition*), threat *reversal* (1^st^ PCA for threat *reversal*) and local activity in the dACC (1^st^ PCA for dACC percent BOLD signal change), respectively. While the overall regression model was not significant for *Bacteroides* (R^2^ = 0.37, F_(25, 33)_= 2.12, *p* = 0.08; **Fig. 4b**) or *Oscillospira* (R^2^ = 0.33, F_(25, 33)_= 1.74, *p* = 0.15, **Fig. 4c**), there were individual variables that were significant. For *Bacteroides*, the model yielded one significant regression weight (β = 0.40, t(25) = −2.80, *p* = 0.01), suggesting an association with connectivity responses during threat *acquisition* (represented by 1^st^ PCA for threat *acquisition*). *Oscillospira* abundance was significantly associated with local activity in the dACC (1^st^ PCA for dACC percent BOLD signal change) (β = 0.52, t(25) = −2.81, *p* = 0.01). Variance in *Prevotella* (R^2^ = 0.25, F(25, 33)= 1.16, *p* = 0.36; **Fig. 4d**) was not related to any specific set of brain measures. Multiple regression results are reported in full in Supplementary Table 5.

**Fig. 4.**
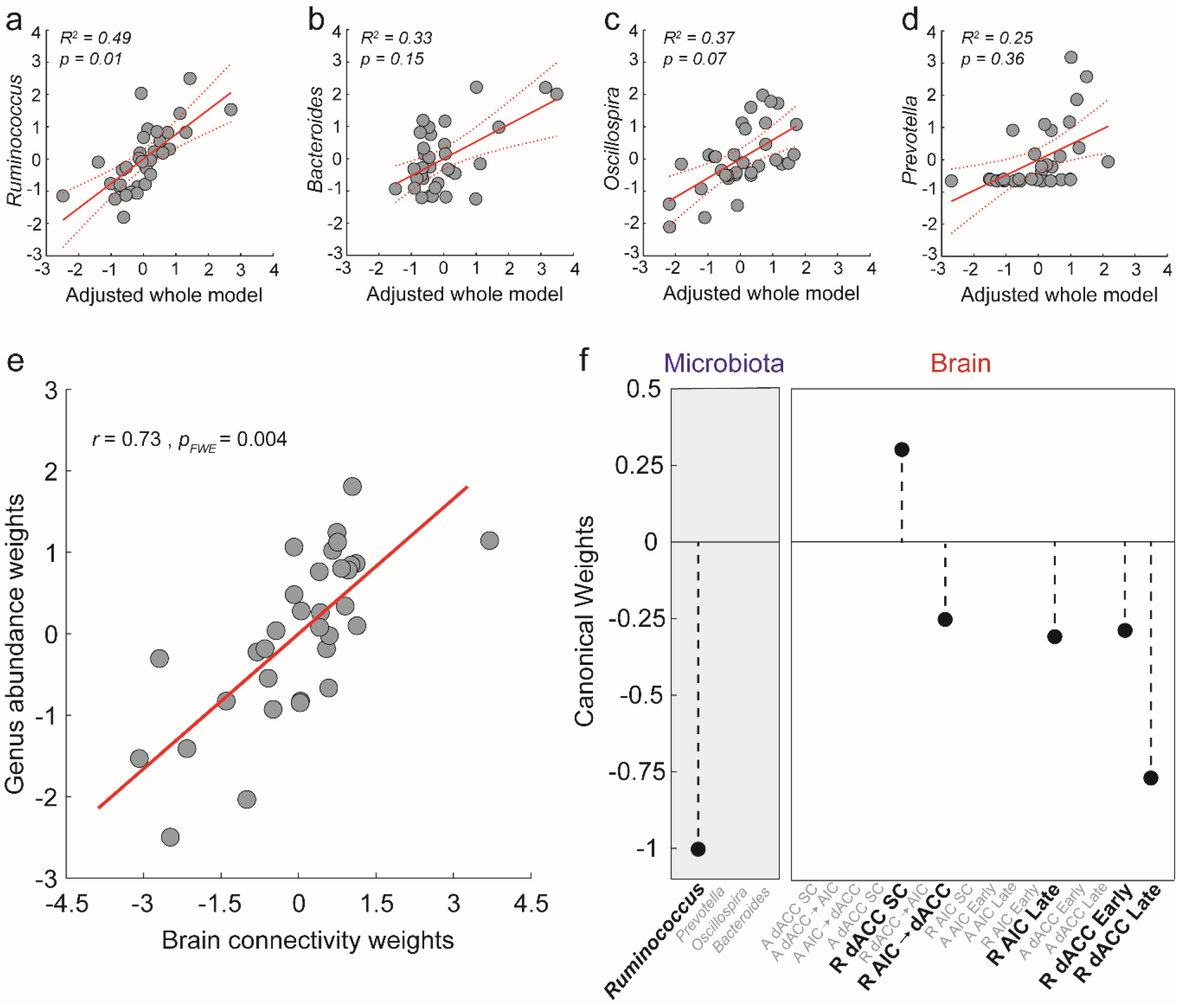
Associations between the driving genera and threat-related brain processes. Principal components analysis (PCA) was used to reduce the dimensionality of brain variables, which resulted in 7 PCs representing connectivity strengths during threat learning (task acquisition phase), threat reversal (task reversal phase), and percent BOLD signal change responses at the AIC and dACC. Results from four independent multiple regressions showed that brain responses predicted the relative abundance of *Ruminococcus* (**a**). While there were individual regression weights that were significant for (**b**) *Bacteroides* and (**c**) *Oscillospira*, the overall regression model was not significant. Variance in (**d**) *Prevotella* was not related to any specific set of brain measures. 95% confidence intervals are represented by the dashed red lines. (**e**) Multivariate analysis (sparse CCA, sCCA) showed a single significant mode of population covariation linking threat processing measures of brain activity and effective connectivity with *Ruminococcus* abundance. (**f**) Bold text shows microbiome and brain weights (coefficients) contributing to the sCCA. Features in grey text represent zero-contributing features to the sCCA, as imposed by the **l_1_**-norm penalty term. Brain variables prefixed with an ‘A’ refer to those occurring in the task acquisition phase, ‘R’ refers to brain variables in the task reversal phase, and ‘SC’ refers to modulatory self-connections.

### 3.7 Multivariate association between driving genera and threat processing brain dynamics

We next investigated the hypothesis that the microbiota and AIC-dACC activity exhibit a multivariate relationship – i.e. multiple dimensions along the genus axes could map to multivariate brain patterns. This hypothesis is motivated by the fact that driving genera are unlikely to exert independent effects on the neural substrates of threat processing. Results from a sparse canonical correlation analysis (sCCA) showed a single mode of population variation linking threat-related brain responses with the driving genera (**Fig. 4e**, *r* = 0.73, *p_FWE_ =* 0.004, see Methods). This analysis identified a significant canonical mode that associated increased Ruminococcus abundance with stronger feedforward connectivity from the AIC to dACC, and local activity within the AIC and dACC during threat reversal (**Fig. 4f**). The stability of the significant sCCA was supported by a leave-one-out (LOO) cross-validation analysis. This analysis involved performing 32 sCCAs using the same sparsity parameter as the original sCCA, but with each iteration removing one subject (r = 0.73 ± 0.01, 0.69-0.77 [mean ± SD, range], Supplementary Figure 5). To further confirm the stability of the sCCA, we performed a secondary cross-validation by randomly removing 15% of the dataset (permutations = 1000) and assessing the stability of the CCA correlation and weights (r = 0.74 ± 0.03, 0.61-0.82, [mean ± SD, range], Supplementary Figure 5b).

### 3.8 Potential mechanisms linking genus abundance to threat-related brain processes

Growing preclinical evidence supports a causal relationship between the production of short-chain fatty acids (SCFAs) - including butyrate, propionate, and acetate - and host behaviour (Stafford et al., 2012; van de Wouw et al., 2018). Specifically, it has recently been suggested that SCFAs may be critical modulators of neuronal functions associated with threat *reversal* (Stafford et al., 2012) and anxiolytic effects (Burokas et al., 2017). In the light of these findings, we assessed whether the driving genera linked to threat-induced AIC-dACC network activity were major contributors to the microbial production of SCFAs. Given that microorganisms work in consort to perform and maintain metabolic functions (c.f. cross-feeding relationships (Baxter et al., 2019; Belenguer et al., 2006)), we started by assessing how the driving genera interact with the broader microbiota ecosystem. To achieve this goal, we constructed interaction networks highlighting co-abundance (positive correlations, r > 0.39 and co-exclusion (negative correlations, r < −0.39) relationships between the driving genera and the wider microbial community (**Fig. 5a-c**). The co-abundance networks revealed that nodes within the *Oscillospira/Ruminococcus* network (**Fig. 5a**) were negatively correlated with nodes within the *Bacteroides* network (**Fig. 5b**). A confirmatory network analysis using *Sparcc* correlations(Friedman and Alm, 2012) supported the above results (Supplementary Figure 6).

**Fig 5.**
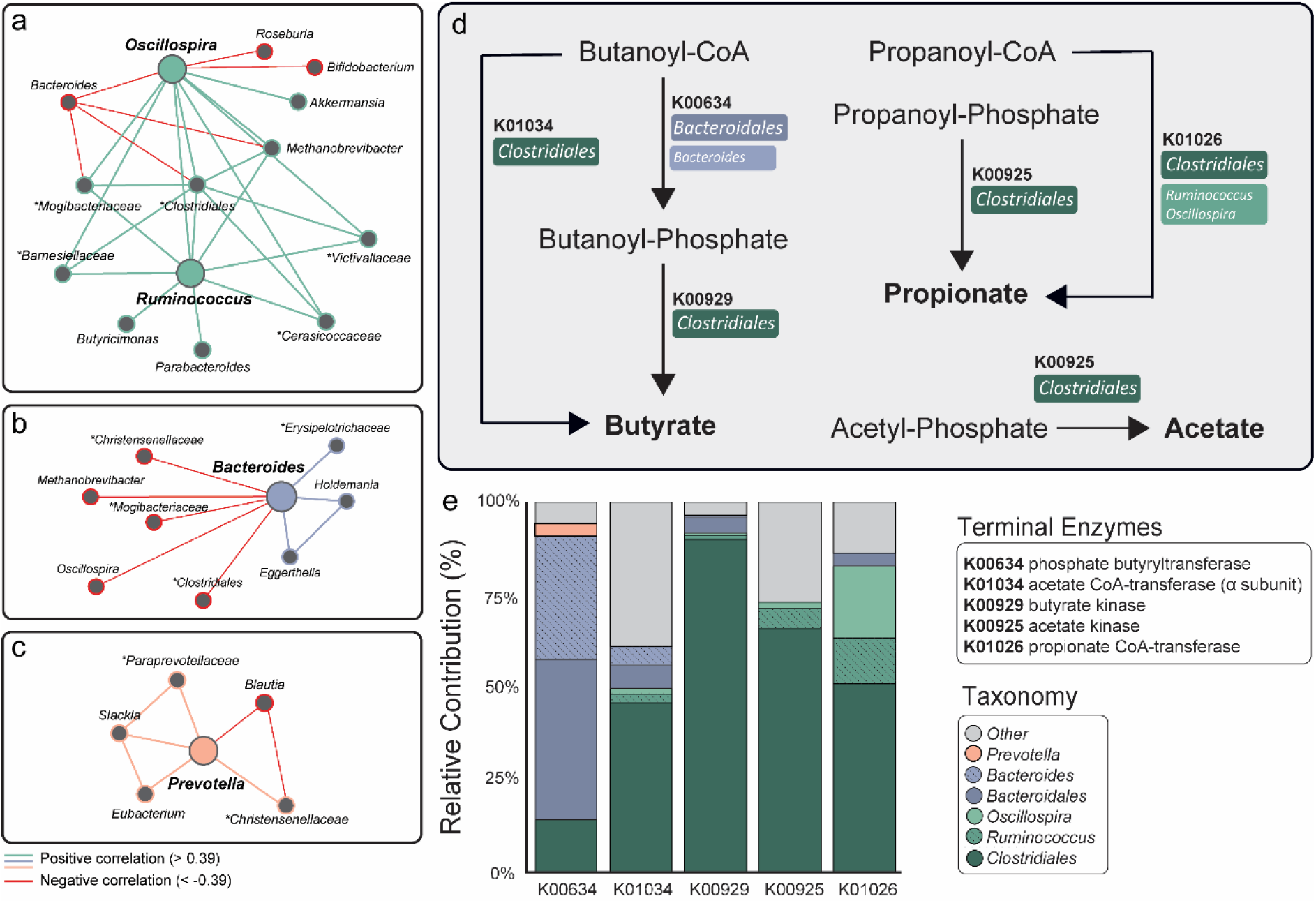
Link between microbiota genera associated to threat-related brain processes and functional pathways supporting the production of short-chain fatty acids. Co-occurrence and co-exclusion relationships between the driving genera (large nodes) including (**a**) *Ruminococcus/ Oscillospira,* (**b**) *Bacteroides,* and (**c**) *Prevotella.* Unclassified genera are described at a broader taxonomic rank above genus level (i.e., family or order) and are marked by asterisks. Graphs are visualized as a force-directed layout using Gephi (Version 0.9.2), using the force atlas template(Bastian et al., 2009). (**d**) Metabolic pathways (derived from the Kyoto Encyclopaedia of Genes and Genomes pathways) representing final enzymatic conversions (terminal enzymes) involved in butyrate, propionate, and acetate production. The major contributor(s) to each gene-encoding enzyme have been identified in colored boxes. (**e**) Decomposition of core/major genera and orders contributing to SCFA pathways. Dark tones represent contributions from a higher taxonomic rank: order. Hatched and lighter tones represent contributions from driving genera.

We then predicted the functional capacity of each microbial sample from its 16S profile using PICRUSt (Phylogenetic Investigation of Communities by Reconstruction of Unobserved States (Langille et al., 2013)). This algorithm predicted the functional capacity of microorganisms by using ancestral-state reconstruction to estimate which gene families are present and how they interact to form the composite metagenome. Functional predictions were then mapped onto the Kyoto Encyclopaedia of Genes and Genomes (KEGG) (Kanehisa and Goto, 2000) to isolate the gene content encoding for terminal enzymes involved in the production of SCFAs (i.e., butyrate, propionate, and acetate) (**Fig. 5d**). The estimated contributions (i.e., the degree to which each microorganism contributed to each sample’s gene content) were represented at both the genus and order scale, and were summarized at the group level (**Fig. 5e**). Members of the *Clostridiales* order, including *Ruminococcus* and *Oscillospira,* were identified as major contributors to all terminal enzymes involved in SCFA production. Other genera within the *Clostridiales* order, including *Blautia, Lachnospira,* and *Coprococcus* were also contributors to these pathways. *Bacteroidales* (order), with a substantial contribution from *Bacteroides* (genus) (**Fig. 5e**), were involved in the sub-terminal reaction (K00634) in butyrate production. As showed in **Fig.4e**, *Ruminococcus* is the strongest multivariate driver of brain patterns. Crucially, results from our functional analysis highlight that the SCFA production pathways also rely on *Ruminococcus* and the order (*Clostridiales*) from which it originates. These findings are consistent with the notion that the detected microbiota features contributing to gut-brain associations supporting threat processing are linked to SCFA production, either directly or in consort with co-occurrence interactions.

## 4. Discussion

We used a multidisciplinary approach to assess the relationship between the human gut microbiota and a brain circuit supporting the subjective experience of threat processing via cognitive-interoceptive appraisal mechanisms (Fullana et al., 2016; Harrison et al., 2015; Kalisch and Gerlicher, 2014). Results highlighted an association between microbial abundance patterns - reflected across different taxonomic scales - with neural dynamics between the AIC and dACC during threat learning and threat updating processes. By considering broad ecological relationships and estimated functional properties of the microbiome, we provide support for the notion that microbial genera influencing threat-related brain processes are involved in SCFA production. More broadly, our findings provide impetus to pursue future research assessing the viability of the gut microbiome to impact brain activity and behaviour linked to threat-related disorders (Harrison et al., 2015).

High-level ecological measures are thought to recapitulate key organizational principles of the gut microbiota (Ley et al., 2006; Lozupone et al., 2012; Mariat et al., 2009). We found that inter-individual variability in α-diversity - a joint measure of the evenness and richness of residing microorganisms - was associated with the strength of inhibitory patterns of connectivity from the dACC to the AIC, and self-inhibitory responses within the AIC during threat *reversal* (**Fig. 3e-f**). The reversal of learned threat stimuli associations requires an updating process, where former safety signals are re-evaluated as a new threat. The α-diversity-mediated modulation of dACC to AIC is consistent with previous work, suggesting that processes involving threat updating are supported by cognitive computations within the dACC (Savage et al., 2020; Stevens et al., 2011). These findings are noteworthy, as α-diversity is thought to engender a microbial ecosystem that is robust and resilient to environmental perturbation (Bokulich et al., 2016). Higher functional redundancy allows the microbiota to compensate for the functions of absent species, including adequate production of microbiota-derived metabolites to meet host demands (Valdes et al., 2018). Accordingly, reduced α-diversity has been linked to exacerbated fear reactivity in infants (Aatsinki et al., 2019; Gao et al., 2019) and altered insular connectivity in adults (Curtis et al., 2019). The Bacteroidetes to Firmicutes ratio was not linked to threat-related AIC-dACC patterns of activity and connectivity. While some evidence suggests a link between this ratio and inflammation (Verdam et al., 2013), obesity (Ley et al., 2006), and chronic pain (Labus et al., 2017), the utility of phyla-level ratios as a biomarker in mental health remains debated. Findings from the current study do not support the putative link between the Bacteroidetes to Firmicutes ratio and threat-related brain processing.

While diversity measures are important from a whole systems approach, when considered in isolation it is unclear whether effects are driven by highly conserved features (i.e., common to all members of major phyla), or if they emerge within finer taxonomic scales. The analysis of genus-level composition is thought to represent an intermediate scale; striking a balance between the complexities of interpreting high-level ecological measures, with a reductionist approach that isolates species of interest. Results from the multiple regression and sCCA analysis converge in supporting an association between *Ruminococcus* abundance and activity from the AIC to the dACC, as well as local activity within the AIC and dACC during threat *reversal*. More broadly, *Ruminococcus* appears to link to key brain processes facilitating the update of previously reinforced safety signals to new threat associations (**Fig. 4a&f**).

The biological mechanisms supporting the causal interplay between the gut microbiome and brain processes underpinning human behaviour are not yet fully understood. The microbiota is thought to impact brain activity via the production of SCFAs including acetate, propionate, and butyrate (Dalile et al., 2019). Accordingly, recent preclinical studies suggest that alterations in microbiota-derived metabolites, including SCFAs, contribute to neuronal activity and behaviour linked to threat processing (Stafford et al., 2012; Whittle and Singewald, 2014). SCFAs are signalling molecules that act as histone deacetylase inhibitors (Kratsman et al., 2016; Stilling et al., 2016), as well as endogenous ligands for G-protein coupled receptors, FFAR2 and FFAR3 (Brown et al., 2003). These receptors are expressed on enteroendocrine cells, various immune cells, and vagal afferents (Egerod et al., 2018; Nøhr et al., 2015). While SCFAs have been shown to cross the blood brain barrier (Liu et al., 2015; Sun et al., 2016), this pathway is not thought to be a major route via which SCFAs exert their influence on cortical activity and related behaviour. Enteroendocrine-mediated vagal signalling is considered to be a more direct and accessible route via which SCFAs influence cortical dynamics and behaviour (Bonaz et al., 2018; Lal et al., 2001). Information from vagal afferents converge in the nucleus tractus solitarius, which can then be relayed to cortical brain regions including the AIC and the dACC. Accordingly, associations between enzymes involved in butyrate production and the insula structure have recently been reported (Labus et al., 2017). Here, we provide evidence indicating that *Ruminococcus* and co-occurring taxa play an important role in the estimated production of acetate, propionate, and butyrate. While our findings are estimations and thus cannot be directly extrapolated, they provide a solid rationale to motivate future work directly testing the putative key role of SCFAs in the modulation of brain processes. Future work that specifically focuses on SCFAs would also support the growing body of existing preclinical and emerging human data (Boehme et al., 2019; Burokas et al., 2017; Dalile et al., 2019; Lee et al., 2020). More specifically, preclinical work suggests a link between *Ruminococcus-*induced increases in SCFAs and improvements in anxiety-related behaviours (Yu et al., 2020), and stress and emotional instability (Provensi et al., 2019). However, important to note is that clinical translation of this work has yielded conflicting results. A recent study in healthy men showed that colonic administration of SCFAs attenuated the cortisol response to physiological stress, but had no effect on threat learning as assessed by subjective ratings and skin conductance responses (Dalile et al., 2020).

A number of caveats need to be considered while interpreting the results from this study. Our analysis focused on a well-defined two-region circuit consistently implicated in human threat processing (Fullana et al., 2016). Moreover, this network has recently been linked to individual differences in anxiety sensitivity – a well validated trait measure of the fear of bodily anxiety sensations (Harrison et al., 2015; Savage et al., 2020). Future work could assess how AIC-dACC circuitry interacts with the broader network of brain regions supporting the contextual processing of threat, and its possible interactions with the microbiota. With regards to the fMRI task, while the use of a fixed ISI has been adopted by previous work (Savage et al., 2020; Schiller et al., 2008), we acknowledge that future work may benefit from a task variant with more events and a non-fixed ISI. We attempted to minimize common confounds associated with both acute (e.g. recent medication use and dietary intake) and general lifestyle changes in microbiota composition by selecting relevant exclusion criteria provided by The Human Microbiome Project Consortium (Methé et al., 2012). Stool sample collection and neuroimaging were also performed in close temporal proximity. However, collection of repeat faecal samples could provide a more nuanced assessment of the relationship between microbiome and brain processes. In the current study, 16S rRNA sequencing was used as this method has been demonstrated to provide sufficient resolution to characterise genus (and broader) level interactions (Rausch et al., 2019). Furthermore, previous work has demonstrated that functional assessments performed using PICRUSt are sufficiently well correlated with genomic content to yield accurate predictions in human gut microbiome samples (Langille et al., 2013). However, as this emerging field continues to develop, the importance of combining neuroimaging and behavioural datasets with higher resolution sequencing (shotgun metagenomics), and untargeted or targeted metabolomics will be critical to extend upon this work. The specificity of SCFAs and their putative mechanistic role in altering AIC-dACC dynamics will also need to be replicated in adequately powered human interventional studies. Current results provide key knowledge and motivation to invest in future targeted work assessing the link between gut features and threat-related neural dynamics.

## 5. Conclusion

There is convincing preclinical evidence demonstrating the effect of the gut microbiota in altering brain mechanisms supporting threat processing. However, there remains a large gap in the understanding of how microbiota consortia engenders variability in neural dynamics underpinning human threat processing. Our study supports distinct interactions between microbial abundance patterns - reflected across different taxonomic scales - with neural dynamics supporting threat learning and threat updating processes in healthy individuals. While research in this field is still in its infancy, current data highlights that the assessment of the microbial milieu may provide insights into the emergence of, and vulnerability to threat-related disorders.

## Author Contributions

Conceptualization and methodology, C.V.H, B.J.H, R.J.M and L.C.; Formal analysis: C.V.H., K.K.I., H.S.S., M.Z., R.J.M., and L.C.; Data curation: C.V.H. and L.A.S.; Resources, G.R.S, L.A.S, L.C.; Writing – original draft, review, and editing, all authors; Funding acquisition, L.C.; Supervision, R.J.M and L.C.

## Acknowledgements

We thank P. Sanz-Leon for her contributions to the fMRI pre-processing pipeline. We thank E. Savage, G. Whybird and N. Koussis for their assistance with data collection, and S. Sonkusare for manuscript feedback. L.C acknowledges support from the Australian National Health and Medical Research Council (NHMRC), APP1099082.

