## Supplementary Material for "Microbiota links to neural dynamics supporting threat processing"

Hall et al.

**Supplementary Note 1**

***Behavioural results supporting successful differential conditioning.*** We used in-scanner ratings (five-point Likert scale, Self-Assessment Manikins) (Bradley and Lang, 1994) to confirm participants' differential subjective aversiveness to the threat (CS+) and safety (CS-) signals at the end of each task phase: *baseline, acquisition,* and *reversal* (**Fig. 1a**). As expected, there were no significant differences in participants' subjective ratings of bodily anxiety sensations (anxious arousal) or unpleasantness (valence) between the CS+ and CS- during *baseline* (where no US was present). We then separately tested the differential response to the stimulus in the *acquisition* and *reversal* phases. Two-factor repeated measures ANOVAs identified a significant main effect of task phase (anxious arousal: $F_{(2,72)}$ = 16.0, $p$ = 1.76 x $10^{-6}$; valence: $F_{(2,72)}$ = 13.1 $p$= 1.37 x $10^{-5}$), stimulus type (anxious arousal: $F_{(1,36)}$ = 34.6, $p$ = 1.01 x $10^{-6}$; valence: $F_{(1,36)}$ = 21.3, $p$ = 4.80 x $10^{-5}$), as well as interactions between task phase and stimulus type (anxious arousal: $F_{(2,72)}$ = 18.5, $p$ = 3.20 x $10^{-7}$; valence: $F_{(2,72)}$ = 15.7, $p_{FWE}$ = 2.22 x $10^{-6}$) (**Fig. 1b**). Post-hoc paired t-tests (Bonferroni corrected) showed significant differences between threat compared to safety signals in the *acquisition* (anxious arousal: $t_{36}$ = 5.99, $p_{FWE}$ = 7.14 x $10^{-7}$; valence: $t_{36}$ = -5.80, $p_{FWE}$ = 1.31 x $10^{-6}$) and *reversal* phases (anxious arousal: $t_{36}$ = 5.68, $p_{FWE}$ = 1.89 x $10^{-6}$; valence: $t_{36}$ = -3.71, $p_{FWE}$ = 6.93 x $10^{-4}$) (**Fig. 1b**). These results confirm that the threat signals evoked greater bodily anxiety sensations and were rated as more unpleasant (negative valence) than the safety signals, during both *acquisition* and *reversal* task phases (**Fig. 1b**). When comparing threat between phases, we observed significant effects for anxious arousal and affective valence when contrasting the *baseline* to the *acquisition* (anxious arousal: $t_{36}$ = -6.25, $p_{FWE}$ = 9.69 x $10^{-7}$; valence: $t_{36}$ =5.56, $p_{FWE}$ = 2.7x$10^{-6}$, respectively), and *baseline* to *reversal* phase (anxious arousal: $t_{36}$ = -5.61, $p_{FWE}$ = 6.96 x $10^{-6}$; valence: $t_{36}$ =5.38, $p_{FWE}$ = 1.40x$10^{-5}$) (**Fig. 1b**). That is, threat signals evoked greater bodily anxiety sensations and were rated as more unpleasant during *acquisition* and *reversal* when compared to *baseline* (**Fig 1a**). Post-hoc paired t-tests showed no significant differences between threat-induced anxious arousal and affective valence in the *acquisition* compared to *reversal* phase.

**Supplementary Note 2**

***Effect of between-region connectivity on local neural response***

To confirm how the observed between-region connectivity affects local neural activity, we tested if *negative* modulation of dACC to AIC connectivity subsequently reduced the local response at the AIC. During threat *reversal*, stronger *negative* modulation of dACC to AIC connectivity was associated with greater inhibitory control within the AIC (R = 0.38, $p$ = 0.03).

This result suggests that signalling from the dACC to the AIC during *reversal* downregulates AIC activity (inhibitory modulation).

**Supplementary Note 3**

**Reducing the dimensionality of the microbiota.** Given the complexity of the gut ecosystem, clustering samples into distinct microbial configurations called enterotypes is proposed to be a useful framework to collapse the high dimensional microbiota into genus features that represent the greatest sources of variability (Arumugam et al., 2011). Using Partitioning Around Medoids (PAM) (Kaufman and Rousseeuw, 2009) clustering, we isolated three 'enterotype' clusters. In line with previous work, between-class analysis revealed that each enterotype was driven by inter-individual variability in the following genera (Falony et al., 2016): *Bacteroides* (enterotype 1), *Ruminococcus/Oscillospira* (enterotype 2), and *Prevotella* (enterotype 3) (Supplementary Figure 2). To confirm the importance of each enterotype-driving genus, we repeated the clustering process three times. At each iteration, we removed the driving genus (both genera for enterotype 2) of each enterotype. Clustering similarity, as assessed by adjusted rank index (where 0 = no overlap, and 1 = identical clustering assignment) (Pinto et al., 2007), showed poor consensus between the original classification compared to when *Bacteroides, Ruminococcus/Oscillospira,* or *Prevotella* were removed (0.31, 0.39, and 0.41, respectively). To confirm that the clusters derived from the enterotype analysis were non-random, we shuffled each sample's relative abundance measures (i.e., shuffling elements within each sample; column) and repeated the enterotype analysis (n = 500 permutations). The Silhouette index of the original dataset compared to surrogate models is significantly higher than chance (0.13 compared to 0.017; Supplementary Figure 3a). The confidence by which the enterotype analysis identified three clusters as the best fit (Calinski-Harabasz index), was significantly higher than the one obtained for the surrogate models (Supplementary Figure 3b).

While the clusters are non-random, a visualization of the data challenged the description of the microbiota in term of three non-overlapping enterotypes (Arumugam et al., 2011). This is because most individuals were distributed along a continuum resulting from log-normal distributions of genus abundance. This is particularly evident for Bacteroides, where the majority of samples occupy an intermediate zone between low- and high-Bacteroides abundance, and thus blurring the boundaries between enterotype 1 and 2 (Supplementary Figure 2). The lack of discrete clusters in the data was further supported by the appraisal of the silhouette index as compared to reference scores (Supplementary Figure 3). An exception to these observations was the Prevotella enterotype, which followed a clear bimodal distribution (i.e., a small fraction of samples had a high relative abundance, while most samples had low or zero abundance). Altogether, these results are in line with a recent shift in the microbiome field, favouring the use of a gradient-based approach to uncover sources of between-subject variation in the microbiota composition (Jeffery et al., 2012).

### Supplementary Figures

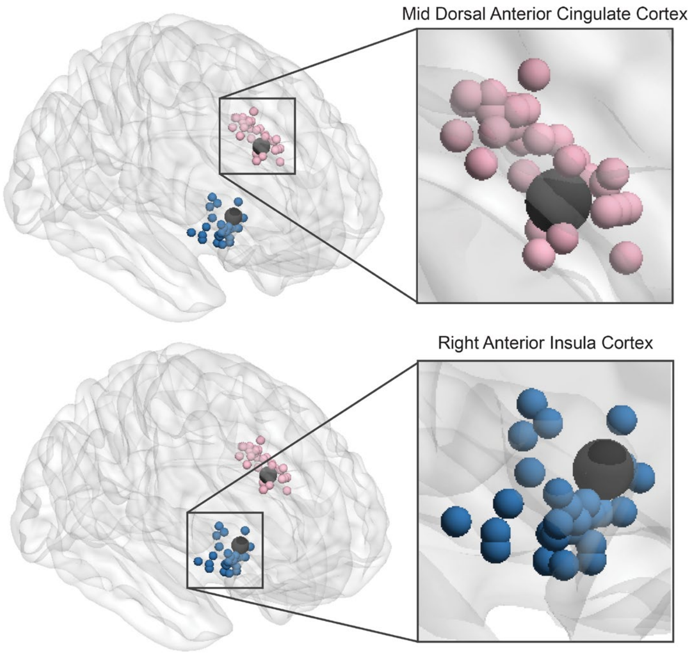


**Supplementary Figure 1. Subject-specific locations of the 5mm regions of interest (ROIs) used to extract time courses at the right AIC (blue) and dACC (pink).** Group-level peak maxima coordinates (obtained from the contrast CS+ > CS-, using 10mm smoothing kernel) used to locate subject-specific ROIs are presented as larger black dots.

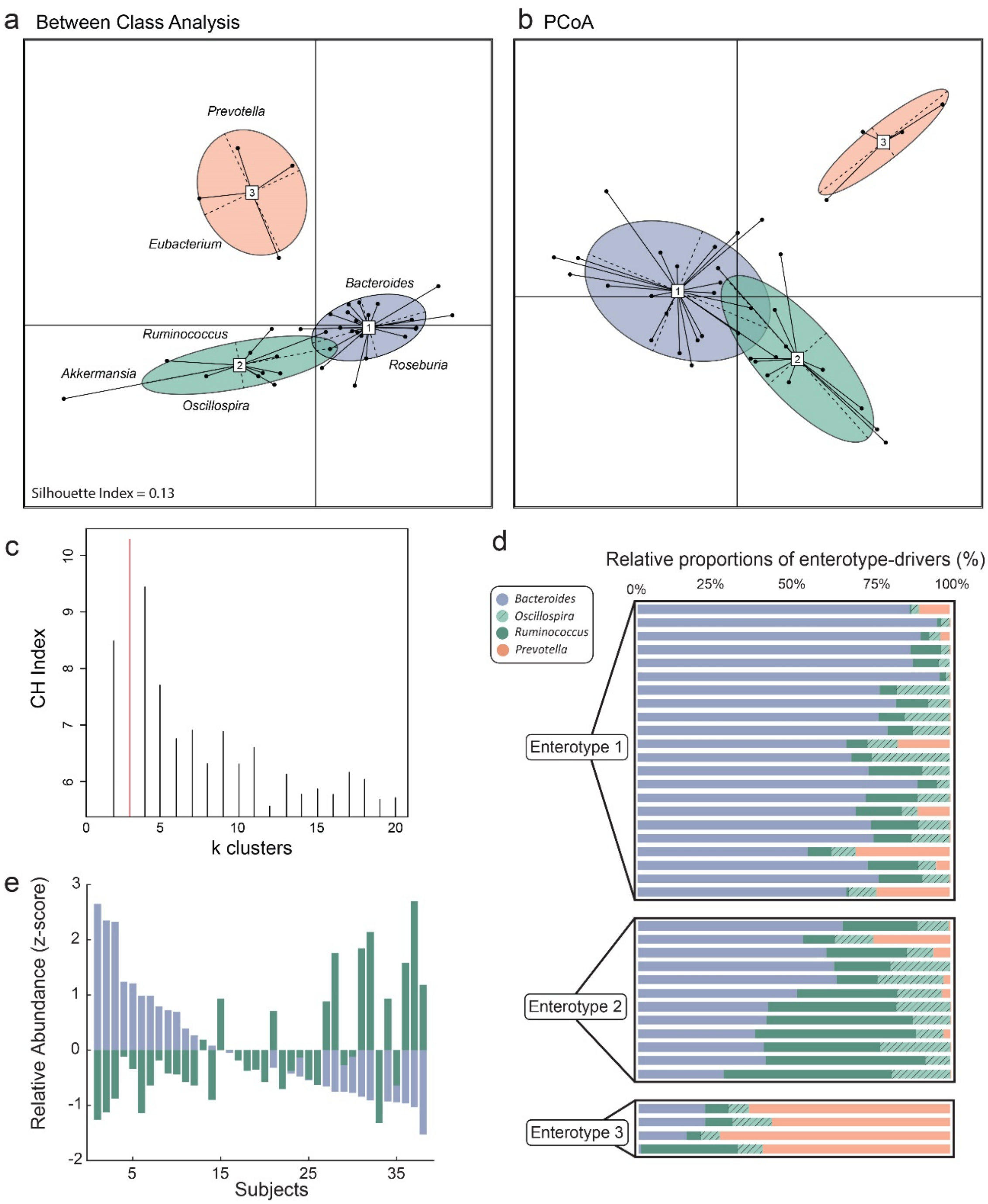


**Supplementary Figure 2. Microbiota-brain interactions within finer taxonomic ranks.** (**a**) Between class analysis and (**b**) PCoA representing three enterotypes, largely determined by inter-individual variability in the relative abundance of select genera, including *Bacteroides* (purple), *Ruminococcus/Oscillospira* (green), and *Prevotella* (orange). However the Silhouette Index, (SI), a measure of how well a sample is matched to its own group, indicated weak evidence (SI = 0.13) for tight enterotype formations. (**c**) The optimal number of clusters was calculated using the Calinski-Harabasz index (Caliński and Harabasz, 1974). (**d**) The relative proportion (%) of each enterotype-driver between the three enterotypes for each individual. (**e**) Subject-specific relative abundances (z-scored) of *Bacteroides* (purple) and *Ruminococcus/Oscillospira* (combined, green), supporting a gradient-based approach over discrete enterotypes (with the exception of *Prevotella*, which formed a bimodal distribution of abundances.

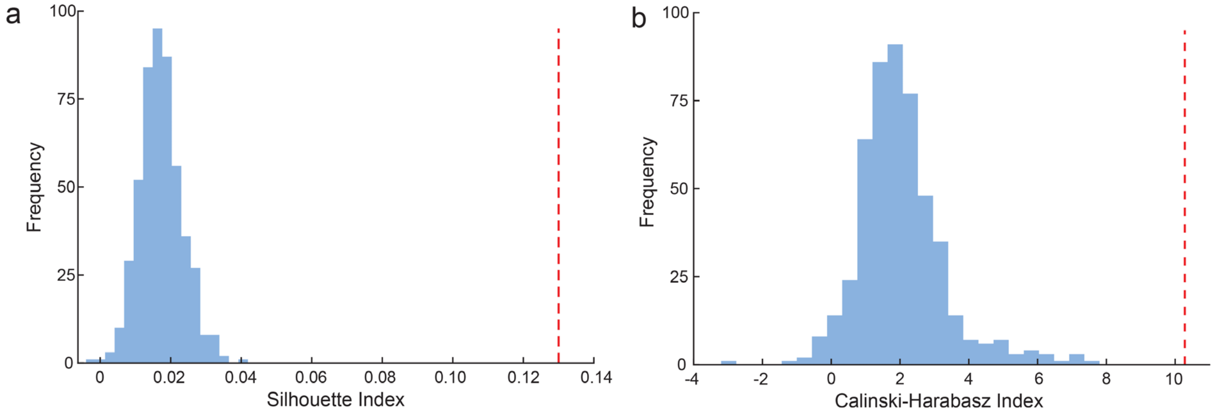


**Supplementary Figure 3. Enterotype control analyses using permutation-based null models.** (**a**) The Silhouette index (a measure of how well a sample is matched to its own cluster) is significantly higher in our dataset (red dashed vertical line), compared to null-based surrogate models where each sample's relative abundances are permuted. (**b**) The Calinski-Harabasz index (a measure of cluster validation where a higher score indicates that clusters are dense and well separated) was significantly higher in our dataset (red dashed vertical line) compared to surrogate models.

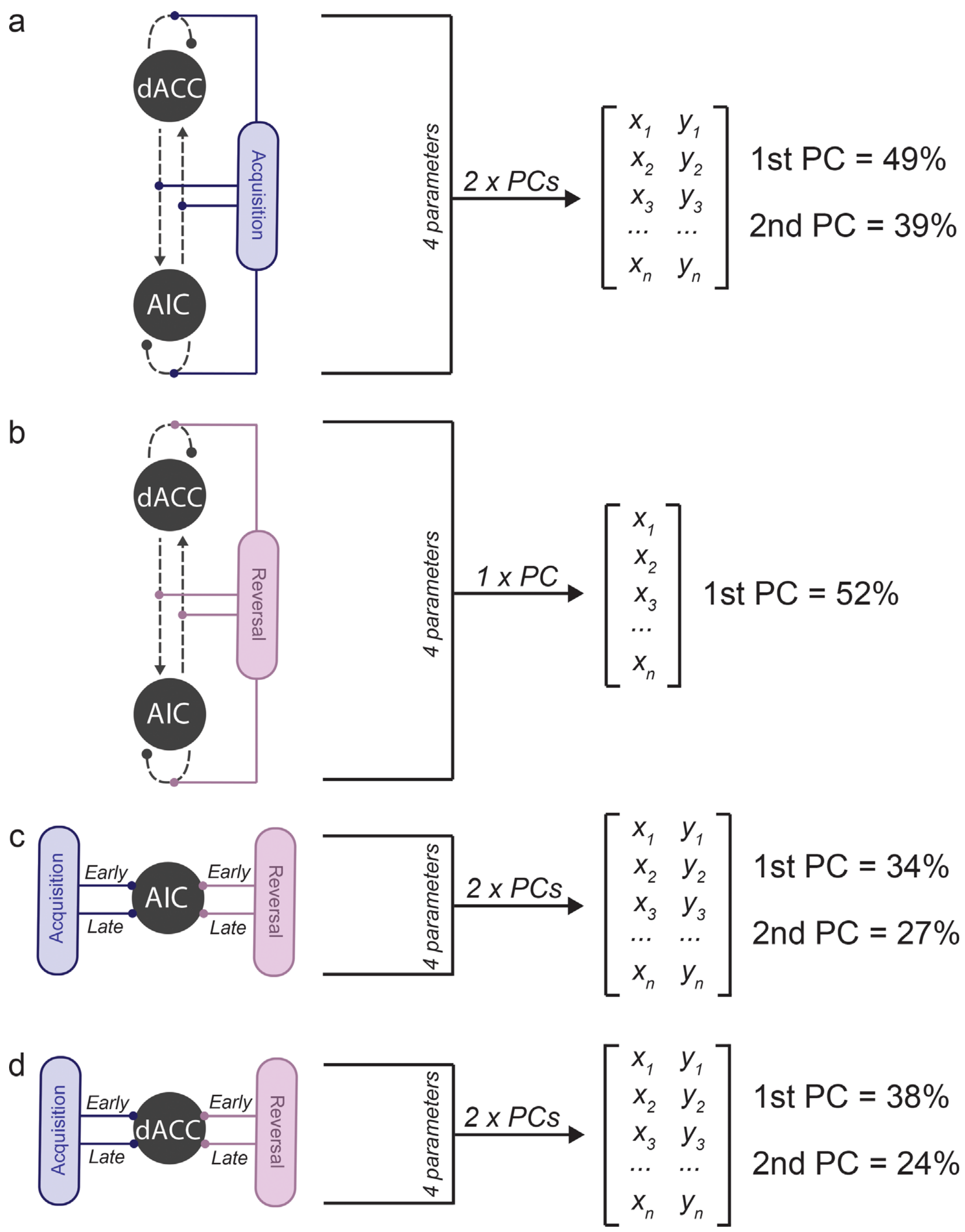


**Supplementary Figure 4. Dimensionality reduction using principal components analysis prior to multivariate analysis.** Multiple principal components analyses (PCAs) were performed to reduce the dimensionality of modulatory connections during (**a**) threat *acquisition* and (**b**) threat *reversal*, each including four parameters each. PCA was also performed on early and late percent BOLD signal change responses at the (c) AIC and (d) dACC, each including four parameters. For threat *acquisition* and BOLD percent signal change responses, the first PC did not exceed 50%. In those cases, a second PC was included. This resulted in 7 PCs used as features in the multiple linear regressions.

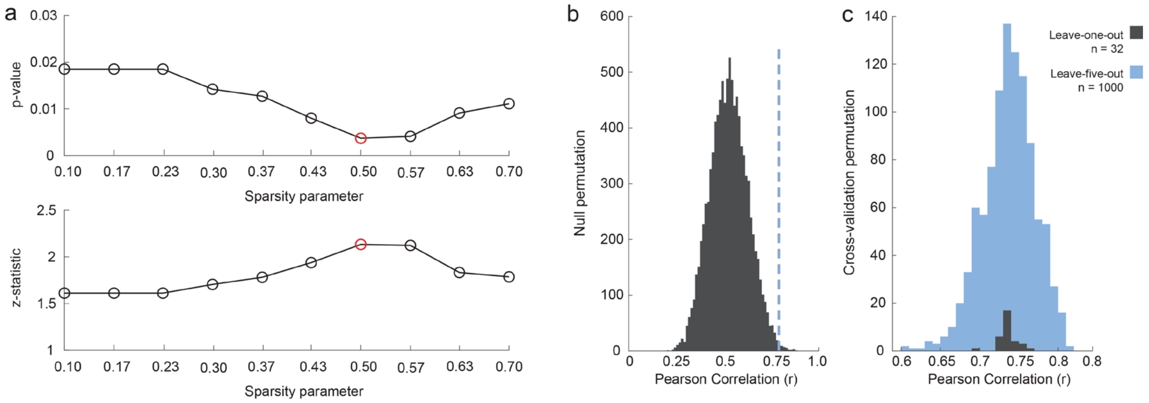


**Supplementary Figure 5. Implementation of the sparse canonical correlation analysis (sCCA).** (**a**) A permutation scheme (n = 10,000) was used to automatically select the best penalty (sparsity) parameter to use for the microbiota and brain variables. Ten default sparsity parameters were tested, ranging from 0.1 – 0.7 (where a lower value indicates a sparser variable space). In line with the p-values (top) and z-statistics (bottom), a penalty of 0.5 was selected as optimal for both microbiota and brain variable sets (red circles). (**b**) Distribution of Pearson correlation coefficients (using the optimal penalty of 0.5) for sCCAs performed on permuted datasets (n = 10,000), compared to the original sCCA correlation (r = 0.74) (dashed, blue line). (**c**) The values of the Pearson correlation coefficients from sCCAs performed on leave-one-out (r = 0.73 ± 0.01, 0.69-0.77, [mean ± SD, range]) and leave-five-out (r = 0.74 ± 0.03, 0.61-0.82, [mean ± SD, range]) datasets were highly consistent with the full dataset (n = 33).

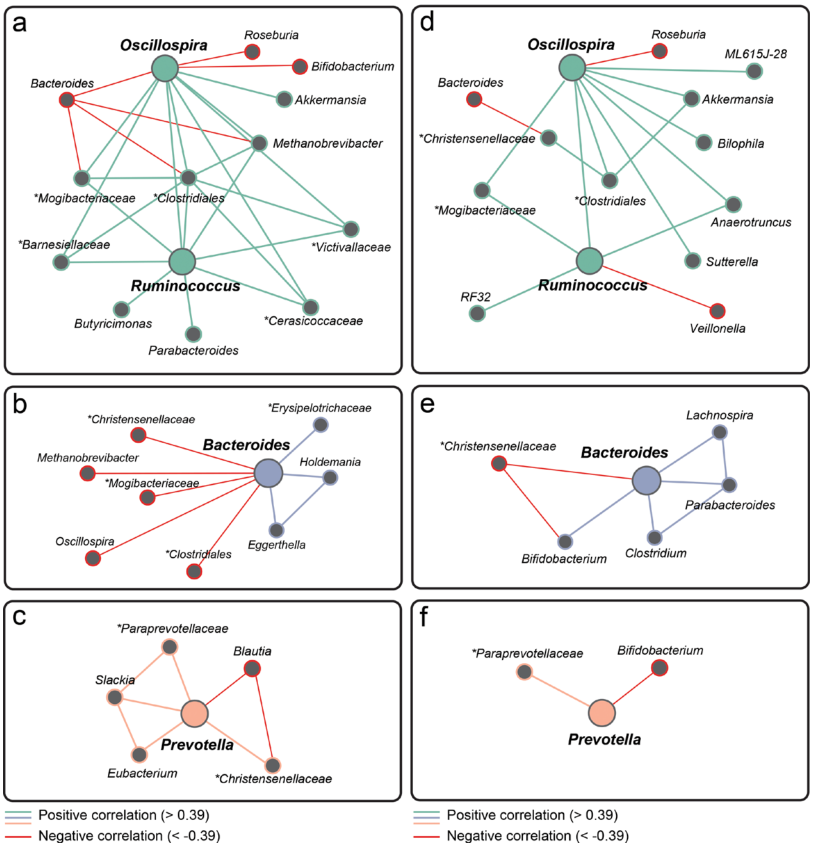


**Supplementary Figure 6.** Comparison of co-occurrence networks for (**a**) *Ruminococcus/Oscillospira*, (**b**) *Bacteroides* and (**c**) *Prevotella* using Pearson correlations on total sum scaled (TSS) and square root transformed data, and (**d-f**) the same driving genera but interactions calculated using *Sparcc* correlations on count data. While the co-occurrence/co-exclusion relationships are more sparse using *Sparcc*, they converge in supporting the existence of a co-occurrence network involving Clostridiales, *Oscillospira*, and *Ruminococcus*, and co-exclusion relationships between *Bacteroides* and genera belonging to the *Oscillospira/Ruminococcus* and *Prevotella* network. Unclassified genera are described at a broader taxonomic rank above genus level (i.e., family or order) and are marked by asterisks. Graphs are visualized as a force-directed layout using Gephi (Version 0.9.2), using the force atlas template.

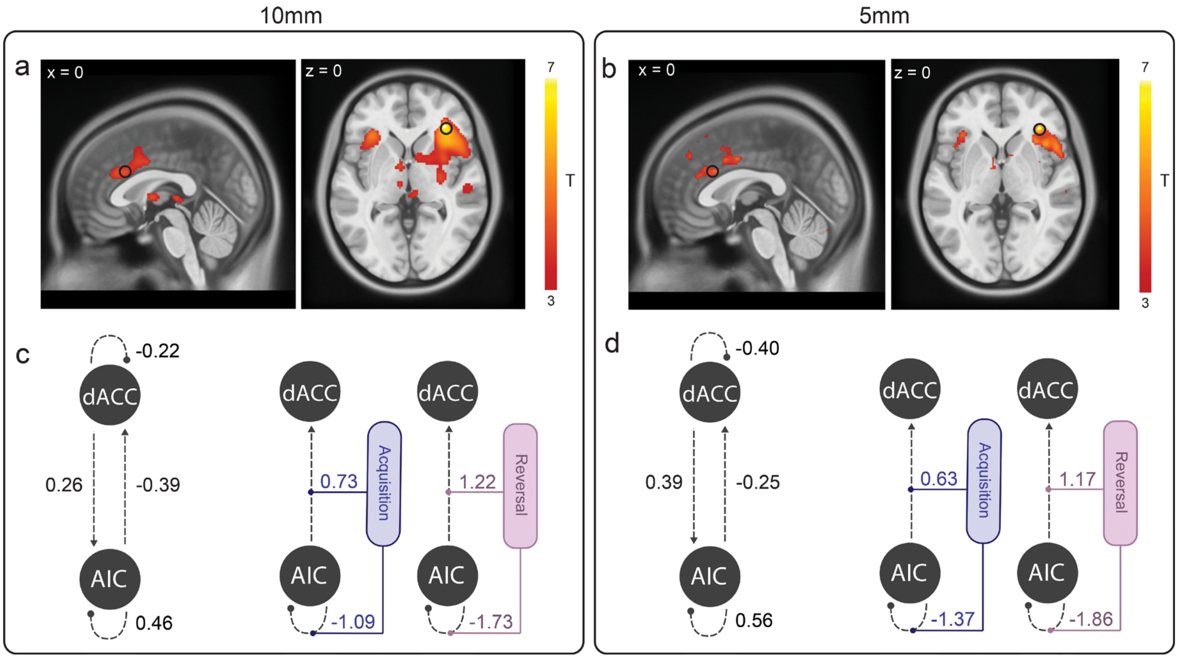


**Supplementary Figure 7. Effect of smoothing kernel on neuroimaging results.** Our neuroimaging results were replicated using a smoothing kernel of 5mm. (**a-b**) Results of the second-level GLM show consistent group-level activation of the dACC and right AIC with the same peak maxima coordinates and similar effect sizes between both analyses. (**c-d**) Three DCMs were re-estimated for each subject using a 5mm smoothing kernel. Bayesian model selection identified the first model (threat-based signals) as the best fit (exceedance probability of 0.6), consistent with the original analysis (Fig. 2b). Finally, we examined the group-level effect sizes of the effective connectivity parameters, and observed similar responses for the endogenous (A-matrix) and modulatory (B-matrix) connections between the 5mm and 10mm smoothing kernels.

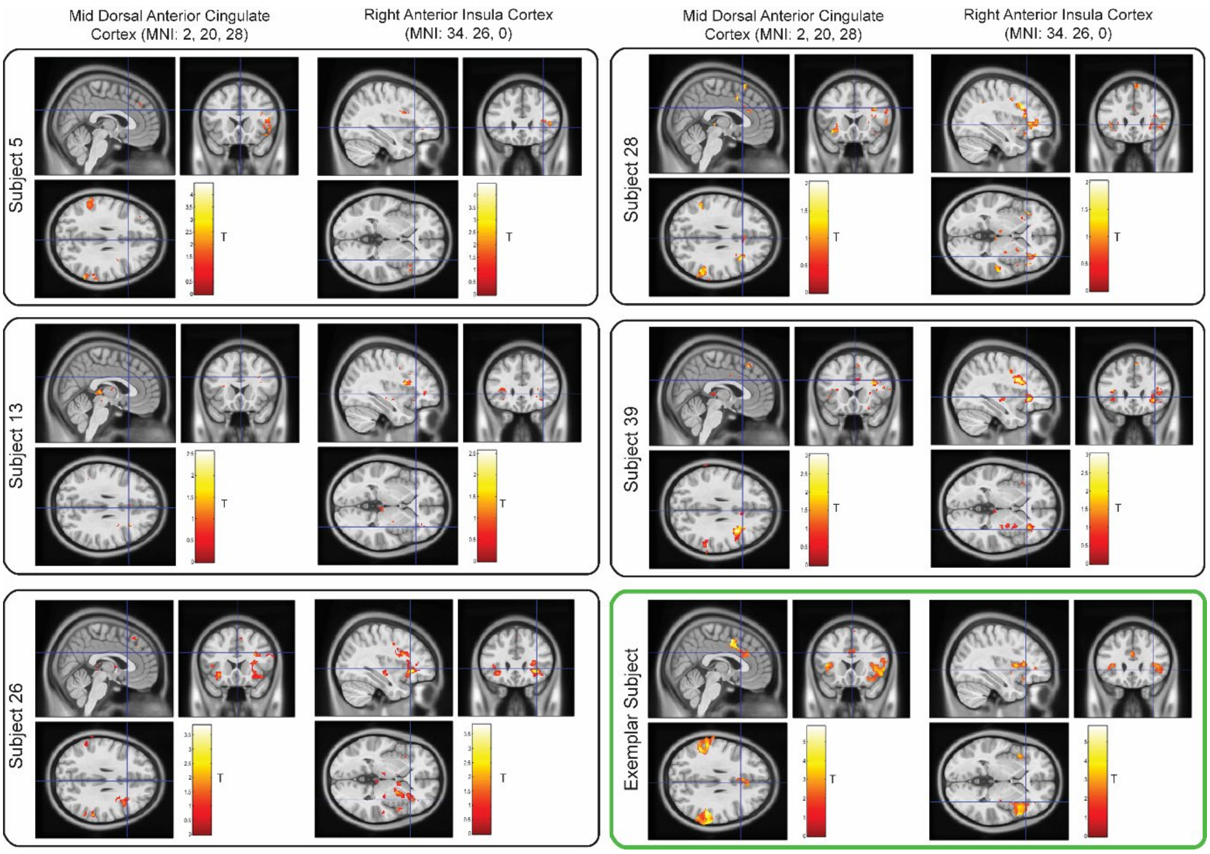


**Supplementary Figure 8. Participants excluded from DCM, PEB, and multivariate assessments.** Upon inspection of subject-specific dACC and AIC fMRI responses, we excluded five subjects who did not exhibit a sufficient response at these two brain regions or where model inversion failed to converge (subjects 5, 13, 26, 28 and 39). The bottom right panel (green) shows an exemplar subject, showing robust single-subject activations at both brain regions ($p_{uncorr.} < 0.05$). Inspection of the pre-processed outputs and nuisance regressors for the five excluded subjects did not highlight any specific problem (including distortions, co-registration, normalisation, or head movements). Furthermore, these subjects did not report significantly different in-scanner subjective ratings of anxious arousal or valence compared to the included (responder) subjects (Supplementary Figure 9). Therefore the lack of single-subject fMRI responses is likely due to the fact that the task presented a limited number of time-locked stimuli, a decision made to keep the length of the task reasonable (16 minutes) and maximise participants' engagement.

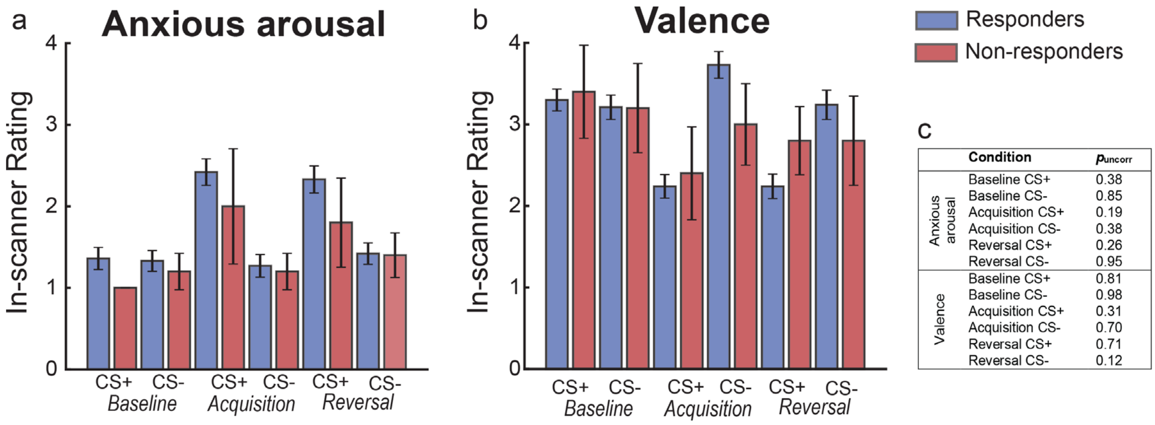


**Supplementary Figure 9.** Non-responder subjects (n = 5) did not exhibit different responses in their subjective ratings of (**a**) anxious arousal or (**b**) valence compared to responder subjects included in the DCM, PEB, and multivariate analyses. (**c**) Independent t-tests ($p_{uncorr.} < 0.05$) confirmed that there were no significant differences observed between any of the subjective in-scanner ratings between the two groups.

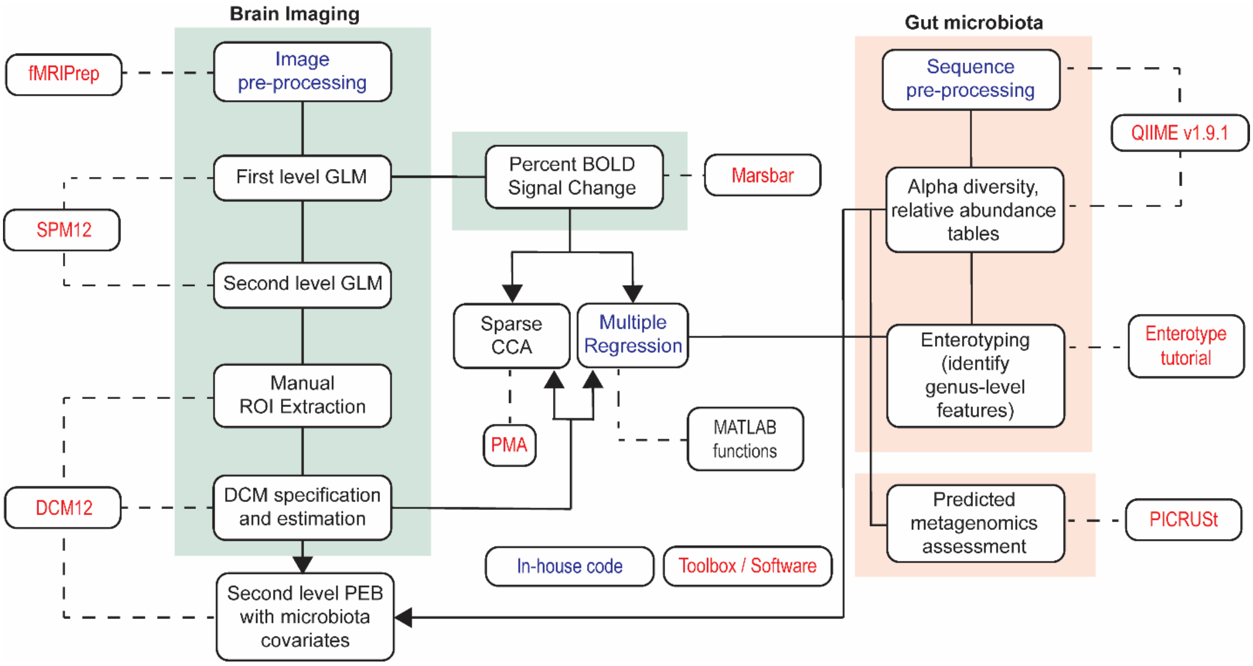


**Supplementary Figure 10. Pre-processing and analysis pipeline for brain imaging (green) and gut microbiota (orange) datasets.** Dashed lines indicate which toolboxes/software/in-house code was required to perform the indicated analysis. Arrows indicate analyses that included combined brain imaging and microbiota datasets.

### Supplementary Tables

#### Supplementary Table 1. Participant Demographics

| Characteristics | |
|---|---|
| Gender | 23 female, 15 male |
| | *Mean ± standard deviation, range* |
| Age, years | 31.7 ± 8.8, 22-50 |
| BMI (kg/m$^2$) | 23.3 ± 2.8, 19.0-29.6 |
| Intelligence Score (WASI-II) | 114.6 ± 11.4, 88-137 |
| Anxiety (HAM-A) | 2.9 ± 3.2, 0-11 |
| Anxiety (HADS) | 5.0 ± 3.6, 0-12 |
| Depression (MADRS) | 2.8 ± 3.6, 0-12 |
| Depression (HADS) | 2.3 ± 2.3, 0-8 |

BMI, Body Mass Index
WASI-II, Wechsler Adult Intelligence Scale
HAM-A, Hamilton and Montgomery Anxiety
HADS, Hospital Anxiety and Depression Scale
MADRS, Montgomery-Åsberg Depression Rating Scale

**Supplementary Table 2.** Comparison of group-level peak maxima coordinates for the right AIC and mid dACC between two different smoothing kernel sizes (5 and 10mm).

| | MNI | | | SPM T | |
|---|---|---|---|---|---|
| **Region** | **x** | **y** | **z** | **5mm** | **10mm** |
| Right AIC | **32** | **34** | **0** | **5.16** | **7.19** |
| Mid dACC | **2** | **20** | **28** | **4.80** | **4.30** |

**Supplementary Table 3. Brain regions activated during threat processing task**

| Cluster size (voxels) | Anatomical region | Peak MNI coordinates | | | |
|---|---|---|---|---|---|
| *Overall threat (CS+) > CS-* | | SPM T-max | x | y | Z |
| **6531** | **R. Anterior insular cortex** | **7.19** | **32** | **34** | **0** |
| | R. Anterior insular cortex | 5.85 | 34 | 26 | 0 |
| | R. Ventral Striatum | 3.73 | 16 | 2 | 2 |
| | R. Superior temporal gyrus | 4.38 | 56 | -22 | -8 |
| | R. Dorsal midbrain | 3.70 | 10 | -14 | -10 |
| **1310** | **R. Dorsal anterior cingulate cortex** | **4.30** | **2** | **20** | **28** |
| | L. Dorsal anterior cingulate cortex | 3.92 | -2 | 10 | 36 |
| | R. Superior frontal gyrus | 3.74 | 10 | -4 | 74 |
| 2253 | R. Supramarginal gyrus (Parietal operculum) | 6.23 | 60 | -42 | 30 |
| 1402 | L. Supramarginal gyrus (Parietal operculum) | 5.64 | -58 | -42 | 26 |
| 1645 | L. Cerebellum | 4.96 | -24 | -64 | -32 |
| | L. Cerebellum | 5.09 | -26 | -74 | -48 |
| 861 | L. Anterior insular cortex | 5.02 | -32 | 26 | -2 |
| 400 | R. Precentral gyrus | 3.69 | 46 | 0 | 56 |
| | R. Middle frontal gyrus | 4.95 | 42 | 0 | 44 |

**Supplementary Table 4. Summary of parameter estimates from Parametric Empirical Bayes (PEB) analysis**

| Endogenous | Group Effects | | | | | |
|---|---|---|---|---|---|---|
| | Ep | Pp | | | | |
| dACC SC | **-0.22** | **1** | | | | |
| dACC → AIC | **0.26** | **1** | | | | |
| AIC → dACC | **-0.39** | **1** | | | | |
| AIC SC | **0.46** | **1** | | | | |
| **Modulatory** | **Group Effects** | | **B/F Ratio** | | **Diversity** | |
| **(*Acquisition*)** | Ep | Pp | Ep | Pp | Ep | Pp |
| dACC SC | -0.37 | 0.89 | 0 | 0 | 0.02 | 0 |
| dACC → AIC | -0.06 | 0.35 | 0 | 0 | -0.16 | 0.54 |
| AIC → dACC | **0.73** | **1** | 0 | 0 | -0.11 | 0 |
| AIC SC | **-1.09** | **1** | 0 | 0 | -0.02 | 0 |
| **Modulatory** | **Group Effects** | | **B/F Ratio** | | **Diversity** | |
| **(*Reversal*)** | Ep | Pp | Ep | Pp | Ep | Pp |
| dACC SC | -0.05 | 0.37 | 0 | 0 | 0.08 | 0 |
| dACC → AIC | 0.00 | 0.33 | 0 | **0** | **-0.34** | **0.97** |
| AIC → dACC | **1.22** | **1** | 0 | **0** | 0.22 | 0.78 |
| AIC SC | **-1.73** | **1** | 0 | **0** | **-0.36** | **0.95** |

Each endogenous (fixed, A-matrix) and modulatory (B-matrix) connection, the mean of the modulatory change in effective connectivity (Ep, expressed in Hz), and the posterior probability (Pp). Significant parameters with a Pp ≥ 95 are indicated in bold.

### Supplementary Table 5. Multiple regression results for each driving genus

| ***Ruminococcus*** | Coefficient Estimate | t-statistic | p-value |
|---|---|---|---|
| Intercept | $-1.54 \times 10^{-15}$ | $-1.10 \times 10^{-14}$ | 1 |
| Threat *acquisition* (PC1) | -0.07 | -0.60 | 0.56 |
| **Threat *acquisition* (PC2)** | **-0.47** | **-2.65** | **0.01** |
| **Threat *reversal* (PC1)** | **-0.37** | **-2.43** | **0.02** |
| AIC percent signal change (PC1) | -0.05 | -0.34 | 0.73 |
| AIC percent signal change (PC2) | 0.06 | 0.34 | 0.74 |
| **dACC percent signal change (PC1)** | **0.45** | **2.71** | **0.01** |
| dACC percent signal change (PC2) | 0.07 | 0.43 | 0.67 |
| ***Bacteroides*** | Coefficient Estimate | t-statistic | p-value |
| Intercept | $-4.45 \times 10^{-16}$ | $-2.75 \times 10^{-15}$ | 1 |
| **Threat *acquisition* (PC1)** | **-0.40** | **-2.80** | **0.01** |
| Threat *acquisition* (PC2) | 0.23 | 1.13 | 0.27 |
| Threat *reversal* (PC1) | 0.11 | 0.63 | 0.54 |
| AIC percent signal change (PC1) | -0.02 | -0.11 | 0.91 |
| AIC percent signal change (PC2) | -0.16 | -0.83 | 0.41 |
| dACC percent signal change (PC1) | 0.05 | 0.24 | 0.81 |
| dACC percent signal change (PC2) | 0.15 | 0.75 | 0.46 |
| ***Oscillospira*** | Coefficient Estimate | t-statistic | p-value |
| Intercept | $3.94 \times 10^{-16}$ | $2.53 \times 10^{-15}$ | 1 |
| Threat *acquisition* (PC1) | 0.08 | 0.59 | 0.56 |
| Threat *acquisition* (PC2) | -0.06 | -0.31 | 0.76 |
| Threat *reversal* (PC1) | 0.06 | 0.35 | 0.73 |
| AIC percent signal change (PC1) | -0.21 | -1.31 | 0.20 |
| AIC percent signal change (PC2) | -0.10 | -0.55 | 0.59 |
| **dACC percent signal change (PC1)** | **0.52** | **2.81** | **0.01** |
| dACC percent signal change (PCA2) | -0.09 | -0.49 | 0.63 |
| ***Prevotella*** | Coefficient Estimate | t-statistic | p-value |
| Intercept | $6.47 \times 10^{-17}$ | $3.78 \times 10^{-16}$ | 1 |
| Threat *acquisition* (PC1) | -0.03 | -0.19 | 0.85 |
| Threat *acquisition* (PC2) | -0.33 | -1.50 | 0.15 |
| Threat *reversal* (PC1) | -0.11 | -0.58 | 0.57 |
| AIC percent signal change (PC1) | 0.24 | 1.39 | 0.18 |
| AIC percent signal change (PC2) | 0.06 | 0.28 | 0.78 |
| dACC percent signal change (PC1) | 0.19 | 0.94 | 0.35 |
| dACC percent signal change (PC2) | -0.10 | -0.47 | 0.64 |

Significant regression coefficients ($p < 0.05$) (excluding the intercept) are indicated in bold.
